# Microbial Electrosynthesis from CO_2_ reaches Productivity of Syngas and Chain Elongation Fermentations

**DOI:** 10.1101/2024.02.08.579422

**Authors:** Oriol Cabau-Peinado, Marijn Winkelhorst, Rozanne Stroek, Roderick de Kat Angelino, Adrie J.J. Straathof, Kunal Masania, Jean Marc Daran, Ludovic Jourdin

## Abstract

Microbial electrosynthesis allows the electrochemical upgrading of CO_2_. However, higher productivities and energy efficiencies are needed to reach a viability that can make the technology transformative. Here we show how a biofilm-based microbial porous cathode in a directed flow-through electrochemical system can continuously reduce CO_2_ to even-chain C2-C6 carboxylic acids during 248 days. We demonstrate a 3-fold higher biofilm concentration, volumetric current density, and productivity than the state of the art, up to a new record of -35 kA m^-3^_cathode_ and 69 kg_C_ m^-3^_cathode_ day^-1^, at 60-97% and 30-35% faradaic and energy efficiencies, respectively. Most notably, the volumetric productivity resembles those achieved in lab-scale and industrial syngas (CO-H_2_-CO_2_) fermentation and chain elongation fermentation. This work highlights key design parameters for efficient electricity-driven microbial CO_2_ reduction. There is need and room to improve the rates of electrode colonization and microbe-specific kinetics to scale-up the technology.

**Graphical abstract:** 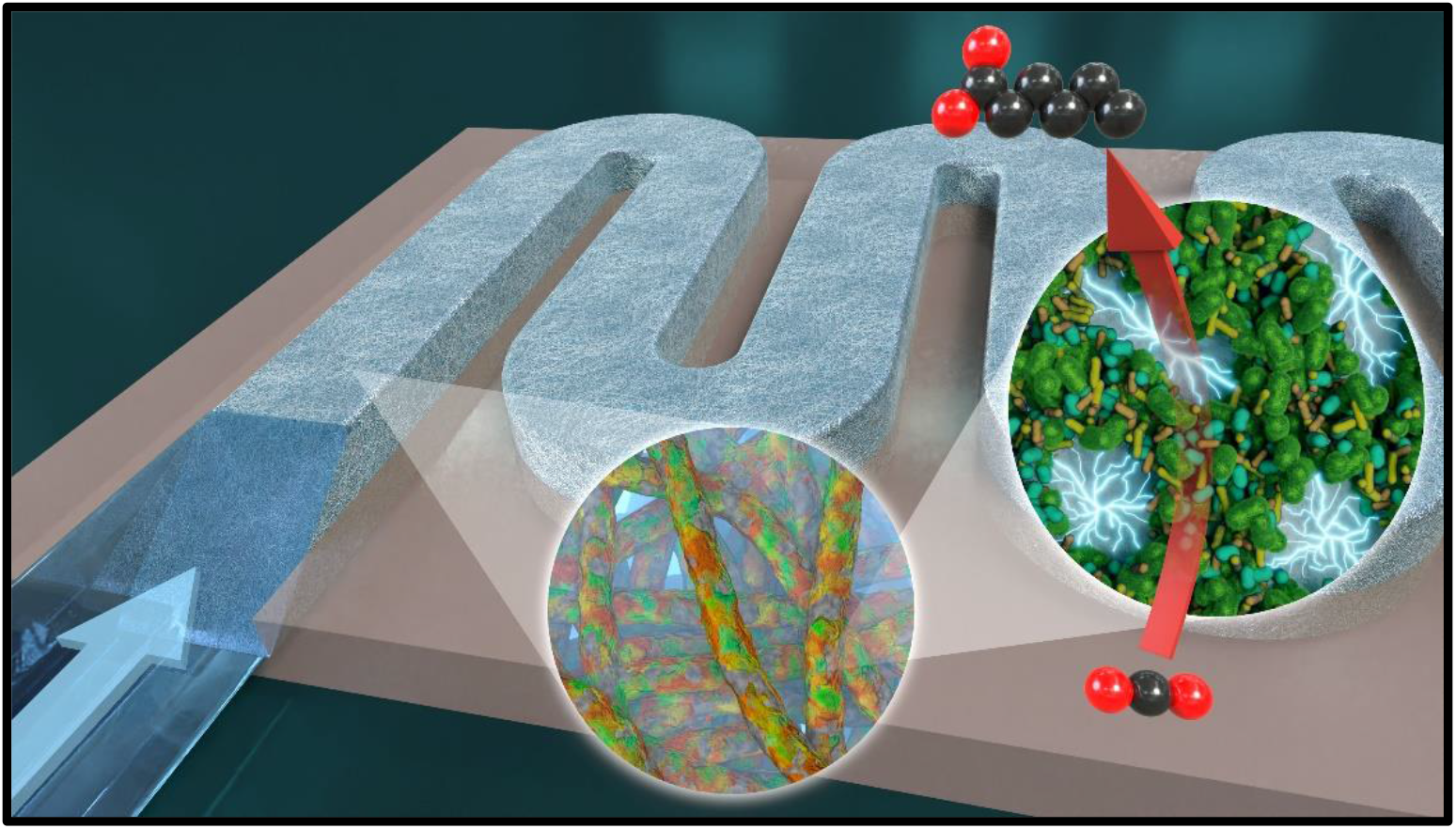

## Introduction

By exploiting the ability of microorganisms to reduce carbon dioxide (CO_2_), microbial electrosynthesis (MES) has become a candidate technology to satisfy the growing demand for fossil-free chemicals synthesis, by harnessing the increasing amount of electrical energy obtained from renewable sources.^1^ Microbiomes are living systems with the ability to self-repair and regenerate, offering a major advantage for resilient industrial applications over abiotic CO_2_ electrolysis. Unlike heterogeneous catalysts, which exhibit stability for limited durations (hours/days), microbial electrochemical reactors demonstrate remarkable operational continuity lasting for several years.^2^ Beyond their robustness, biocatalysts in MES systems exhibit the capacity to generate multi-carbon products with notable selectivity and faradaic efficiency.^3^ The bioelectrochemical reduction of CO_2_ to produce medium-chain carboxylic acids (MCCAs), such as butyric (4 carbons, C4) and caproic (6 carbons, C6) acids, presents a promising avenue for generating low CO_2_ footprint precursors crucial for applications in the fuel, chemical, feed, and food industries.^4, 5^ Nonetheless, it is noteworthy that existing studies elucidating microbial CO_2_ reduction to MCCAs report production rates lower than those achieved in alternative fermentation technologies for organics production, including syngas fermentation and chain elongation fermentation.^6-9^

Following the initial proof-of-concept demonstrating the production of soluble organics from CO_2_ in microbial electrosynthesis,^10^ the primary focus within the MES research community has centered on enhancing microbial catalysts, improving cathode materials, and elucidating fundamental mechanisms and microbial functionalities.^2, 11^ These endeavors have been pivotal in achieving noteworthy key performance indicators (KPIs), including productivities and faradaic efficiencies. Nevertheless, these KPIs have not yet reached a level that ensures the economic viability of the technology.^4, 11^ In the context of transitioning to industrial implementation of MES and its potential role in the electrification of the chemical industry, reactor design emerges as a crucial aspect requiring attention.^2, 11^ State-of-the-art MES reactors capable of producing acids longer than C2 are predominantly biofilm-driven systems, exploiting the proximity to the electron source for CO_2_ reduction.^2, 6, 12^ Biofilm-driven MESs have so far outperformed MES driven by microorganisms in suspension by several orders of magnitude. Nevertheless, biofilms growing in other environments have demonstrated susceptibility to mass transfer limitations, impacting microbial activities due to the necessity for substrates and products to diffuse in and out of biofilms.^13-16^ Despite these limitations, relatively few efforts have been dedicated to researching and developing biofilm-driven MES reactor design concepts that ameliorate mass transport.^6, 17, 18^

The predominant focus in devising innovative reactor designs has been directed towards MES systems employing microorganisms in suspension.^19^ Notably, Cui et al. recently presented an electrolytic bubble column featuring an external hollow fiber membrane gas–liquid contactor for the production of acetate from CO2, achieving an acetate titer of up to 34.5 g L^-1^.^20^ In a different approach, Rosa et al. retrofitted a conventional stirred bioreactor with electrodes, showcasing adaptability to MES applications.^21, 22^ Enzmann et al. introduced a bioelectrochemical bubble-column reactor, serving the dual function of a microbial fuel cell and microbial electrosynthesis system.^23^ Puig et al. explored a tubular MES reactor for the production of butyrate, ^24^ as well as acetate and ethanol at a 1-to-1 ratio.^25^ Additionally, a 4.3 L scaled-up version of a flat-plate double-chamber reactor design demonstrated acetate production by microorganisms in suspension.^26^

Configurations for biofilm-driven microbial electrosynthesis employing flow-through designs, where convective flow is intensified near the biofilm-cathode interface, have resulted in elevated production rates, enhanced biofilm growth, and increased carbon selectivity towards longer MCCAs compared to other tested designs.^6, 13^ However, the flat-plate design employed by Jourdin et al. presents scalability challenges.^6^ In this design, the cathode compartment featured a 1.2 cm-thick free-flowing catholyte volume positioned between the membrane and the 3D-filamentous cathode. The catholyte was directed to flow through the cathode material and exit the compartment on the opposite side, where an additional 1.2 cm-thick free-flowing liquid volume was located. The incorporation of such free-flowing liquid dead-volumes substantially increases both the footprint and capital cost of the reactor upon scale-up. Moreover, these dead-volumes contribute to issues related to hydrogen accumulation, negatively impacting system performance.^27^ The fluid dynamics within this reactor design exhibit suboptimal characteristics, leading to dead zones within the cathode material where mass transport and microbial activity are constrained. Furthermore, the 2.4 cm separation between the anode and the cathode proves to be excessively large, resulting in considerable ohmic and mass transfer resistances, consequently leading to high energy losses.^11^

Here we introduce a directed-flow-through bioelectrochemical reactor (DFBR) featuring a serpentine flow-pattern architecture, as illustrated in Figure 1**A-B**. In this innovative design, CO_2_-saturated catholyte is directed through a continuous serpentine channel entirely filled with a porous 3D carbon-based electrode, where CO_2_ undergoes biological reduction to form medium-chain carboxylic acids (Figure 1**C**). Unlike previously used systems, the DFBR design eliminates free-flowing liquid in the cathode chamber, thereby facilitating substrate and product turnover at the biofilm-cathode surface. Additionally, the serpentine flow-pattern enables an extended residence time, potentially enhancing carbon and electron/hydrogen utilization efficiency, which are key performance indicators upon scale-up. While not explored in this study, the ability to manipulate this residence time theoretically positions this reactor design for further carbon elongation towards carboxylic acids longer than C6. Notably, the design is characterized by its scalability and stackability, enhancing its versatility and applicability.

**Figure 1.**
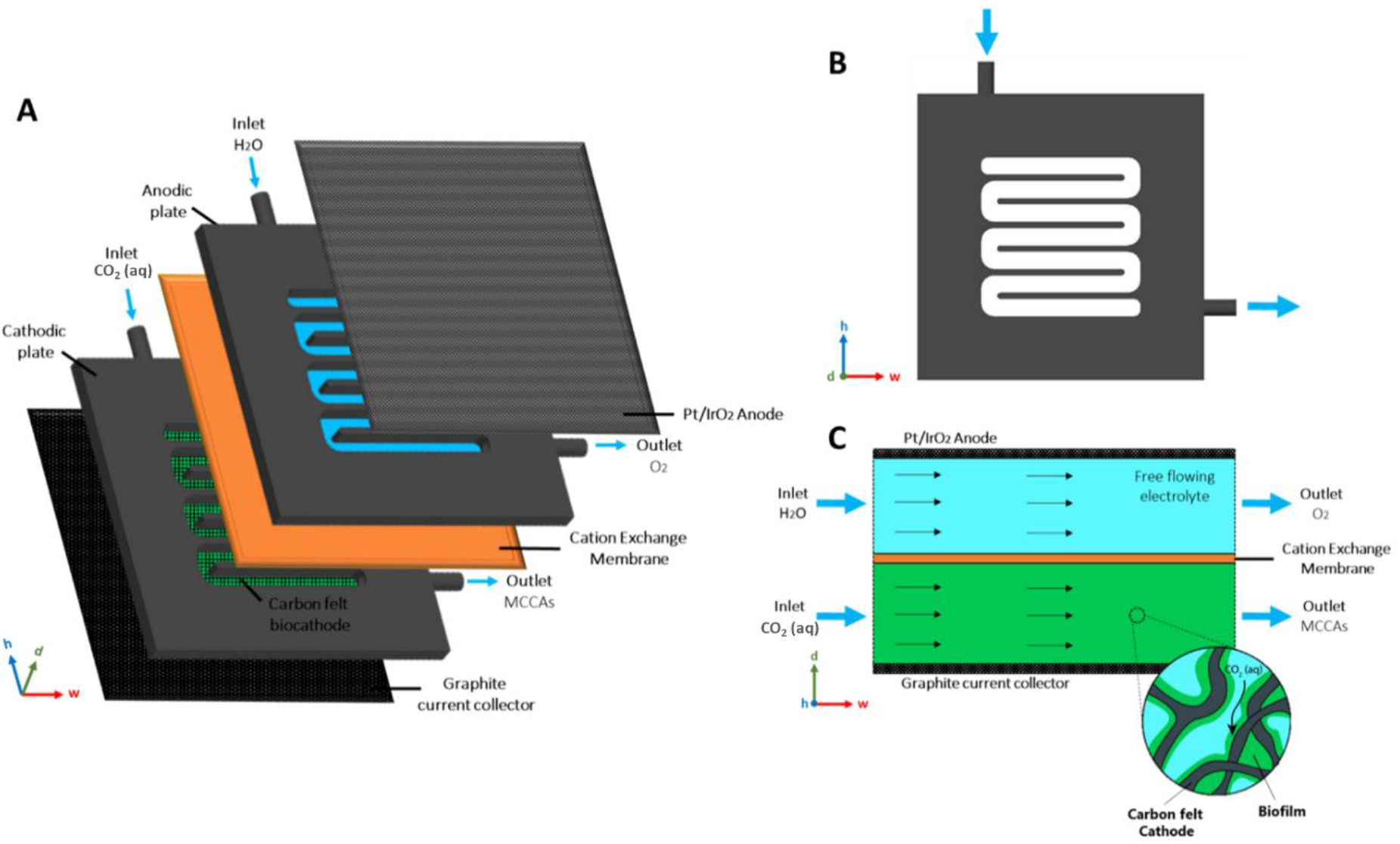
Schematic of a directed-flowthrough serpentine bioelectrochemical reactor. (A) Reactor cell diagram. Each unit consists of a serpentine plate filled with a carbon-based 3D-biocathode on a graphite current collector, an empty serpentine plate (free flowing electrolyte) on a Pt/IrO2 coated 2D-titanium anode, and a cation exchange membrane. (B) Schematic of the serpentine plate design. (C) Top-view schematic of both flow-channels on a reactor cell unit.

The key performance indicators of the DFBR design for the production of MCCAs directly from CO_2_ were investigated. We comprehensively examine the microbial growth, electrode colonization, metabolic activity, organics production, and energy efficiency over an operational period spanning 248 days. 16S rRNA-sequencing allowed to identify the dominant microbial species responsible for the elongation of CO_2_ to MCCAs. Our findings highlight the remarkable capability of microbial electrosynthesis to attain reactor-scale performances comparable to established technologies, establishing the viability of the novel directed-flow-through reactor as a potentially scalable system.

## Results and discussion

In the current investigation, we devised an innovative microbial electrosynthesis reactor, termed a directed-flow-through bioelectrochemical reactor (DFBR). This DFBR introduces a serpentine flow-pattern architecture in both the cathode and anode compartments. At the cathode, the serpentine channel is filled with a carbon felt electrode through which the catholyte is forced to flow through. Two reactors were continuously operated, either potentiostatically (CA) or galvanostatically (CP), for 220 days. Nutrients were replenished using a hydraulic retention time of 4 days, complementing the continuous sparging of CO_2_.

### Biomass growth rate and microbial kinetics of electrode colonization similar in potentiostatic and galvanostatic-controlled reactors

The time-dependent biomass-specific growth rate (*μ*) was experimentally determined for both reactors (**Figure 2A)**, using a recently published method^28^. This method differentiates microbial growth in biofilm and in suspension. The progression of biofilm quantity per electrode volume over time and the associated percentage of electrode colonization under our experimental conditions were assessed (**Figure 2B**). The percentage was calculated on basis of the total biofilm amount, which reached a plateau after 225 days, representing biofilm saturation. This saturation point likely indicates a restriction in space that impeded further biofilm growth.

**Figure 2.**
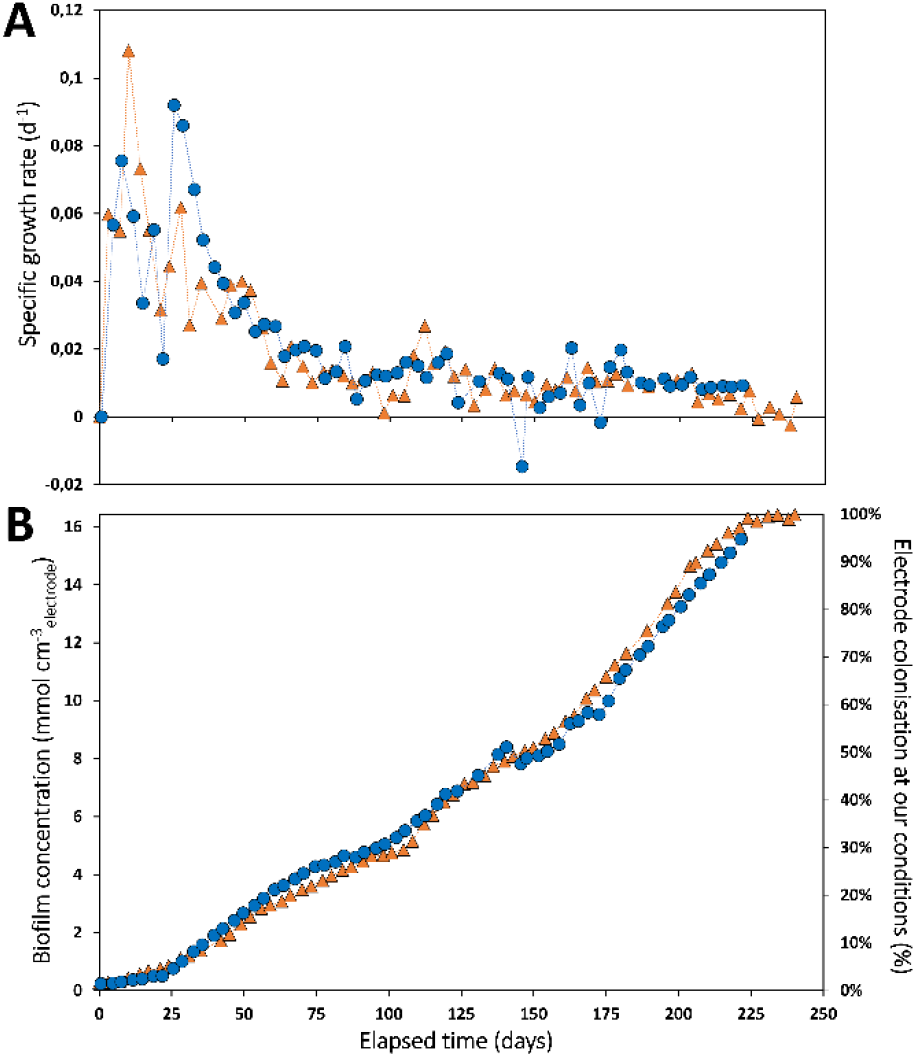
Biomass-specific growth rate (A), biofilm concentration and electrode colonization (B) measured over time in the galvanostatic (orange triangles) and potentiostatic (blue circles) reactors.

In both reactors, growth rates ranging from 0.03 to 0.11 d^-1^ were recorded during the initial 50 days, followed by a decline in growth rates to approximately 0.01 d^-1^ until the conclusion of the experiments. Comparable growth rates and trends were previously observed in non-optimized flow-through MES reactors.^28^ Consequently, the present DFBR did not speed-up biofilm growth and electrode colonization. These growth rates remain modest compared to those observed in analogous anaerobic fermentation technologies like syngas fermentation and chain elongation (up to 2.9-5.7 d^-1^).^29, 30^ Consequently, achieving a fully grown and colonized electrode proved time-consuming in this context (225 days), underscoring the need for improvements from an application perspective. Furthermore, our observations indicated that keeping either current or potential at the used static value did not influence the growth of biofilms and electrode colonization. This suggests that electron uptake from the electrode may not be the limiting process, indicating the presence of other limiting factors influencing growth. This underscores the significant challenge of rapidly colonizing a large electrode, representing a key limitation in the scale-up of microbial electrosynthesis technology.

### Directed-flow-through bioelectrochemical reactor allows three times denser biofilm than previous state-of-the-art

The preceding state-of-the-art in MES ^28^ achieved a biofilm apparent density, or amount of biofilm per electrode volume, of 5.0 ± 2.7 mmol_x_ cm^-3^_cathode_ (Figure S1 in Supplementary Information). In contrast, the novel serpentine design of the directed-flow-through bioelectrochemical reactor remarkably increased the biofilm apparent density (dry cell mass concentration) by over three-fold, reaching 16.4 mmol_x_ cm^-3^_cathode_. Notably, biofilm constituted >99% of the biomass in the reactors (Figure S2), underscoring the DFBR’s efficacy for biofilm-driven bioelectrochemical processes. Attempts to promote biofilm growth in other reactor concepts faced challenges, with microorganisms being washed out during the transition from fed-batch to continuous mode.^31, 32^ A higher biofilm apparent density translates into a larger number of microbes available for the target reaction. As inoculum and electrode material were consistent between our study and Winkelhorst et al. (2023)^28^, this suggests that the reactor architecture and flow pattern/fluid dynamics played a pivotal role in the increased apparent biofilm density. The DFBR design mitigates dead zones, ensuring the entire carbon felt is accessible for biofilm growth. Additionally, the DFBR design eliminates free-flowing liquid in the cathode chamber, facilitating transport of CO_2_, nutrients, H_2_, and products at the biofilm-cathode interface and throughout the channel. It is noteworthy that the catholyte superficial fluid velocity through the electrode in the DFBR was 12 times higher than in Winkelhorst et al. (20.0 vs. 1.7 mm s^-1^), which may have contributed to the observed biofilm density as well. Further investigation is warranted to elucidate the impact of fluid velocity on biofilm in MES. Photographic evidence of the cathodes and SEM images from both reactors at the experiment’s conclusion (Figure 3 and S3) confirmed the development of a thick and dense biofilm, visible to the naked eye, on both sides of the carbon felt, spanning its entire thickness and along the entire serpentine channel. Well-formed biofilm on individual carbon fibres, comprising morphologically diverse microorganisms encapsulated in an extracellular matrix, is observable throughout the entirety of the carbon felt.

**Figure 3.**
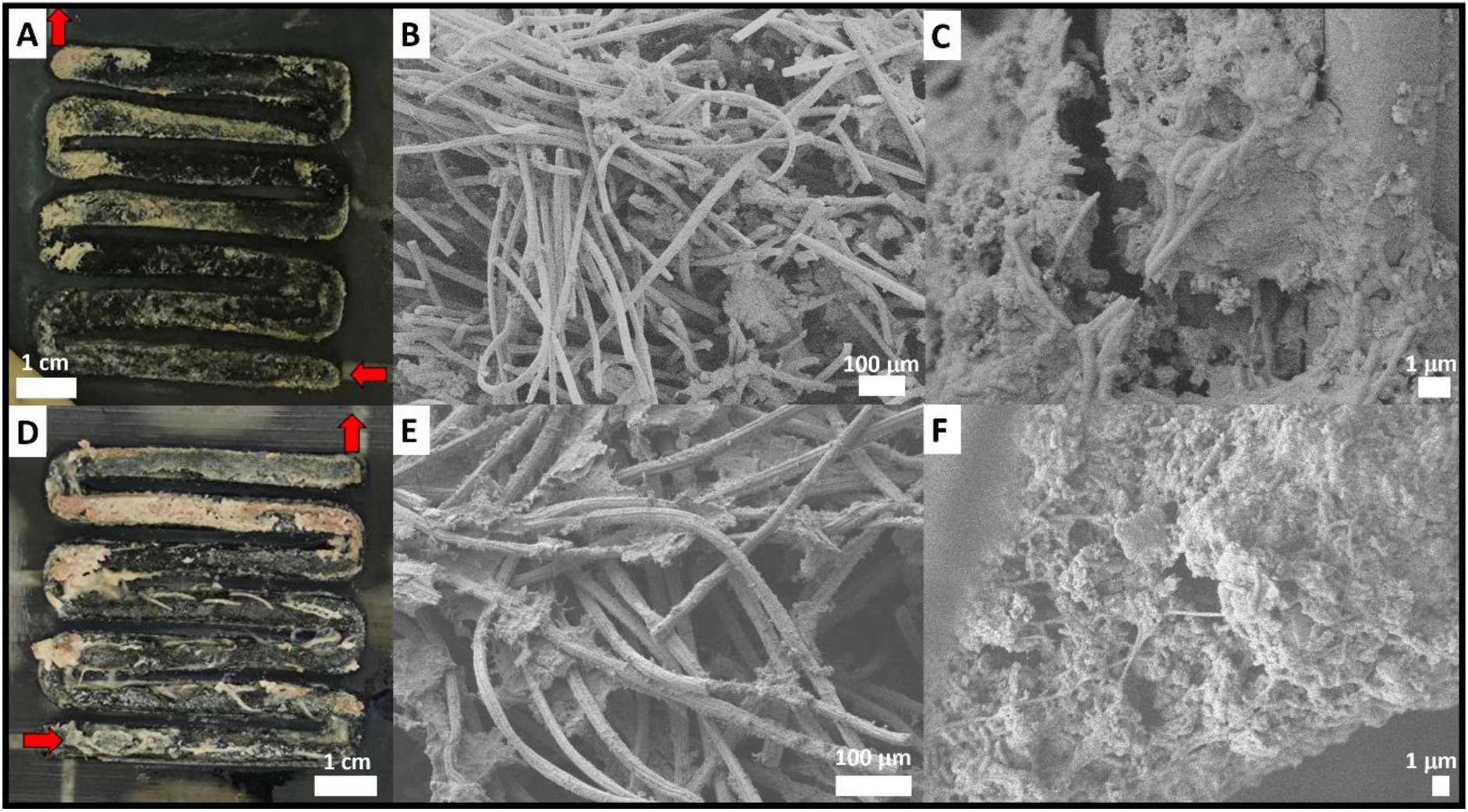
Photograph and scanning electron microscopy images of the biofilm grown on the carbon felt electrode of the galvanostatic (A-C) and potentiostatic (D-F) reactors, at the end of the experiment.

### Directed-flow-through bioelectrochemical reactor results in 5-fold higher volumetric current density and productivity than the state-of-the-art

The evolution over time of the current density, cathode potential, organics concentration, and faradaic efficiency in both reactors were evaluated (**Figure 4**). The trends in organics production rates and current normalized to electrode volume are depicted in Figure S4 and S5. Notably, no alcohols were detected at any point during the experiments.

**Figure 4.**
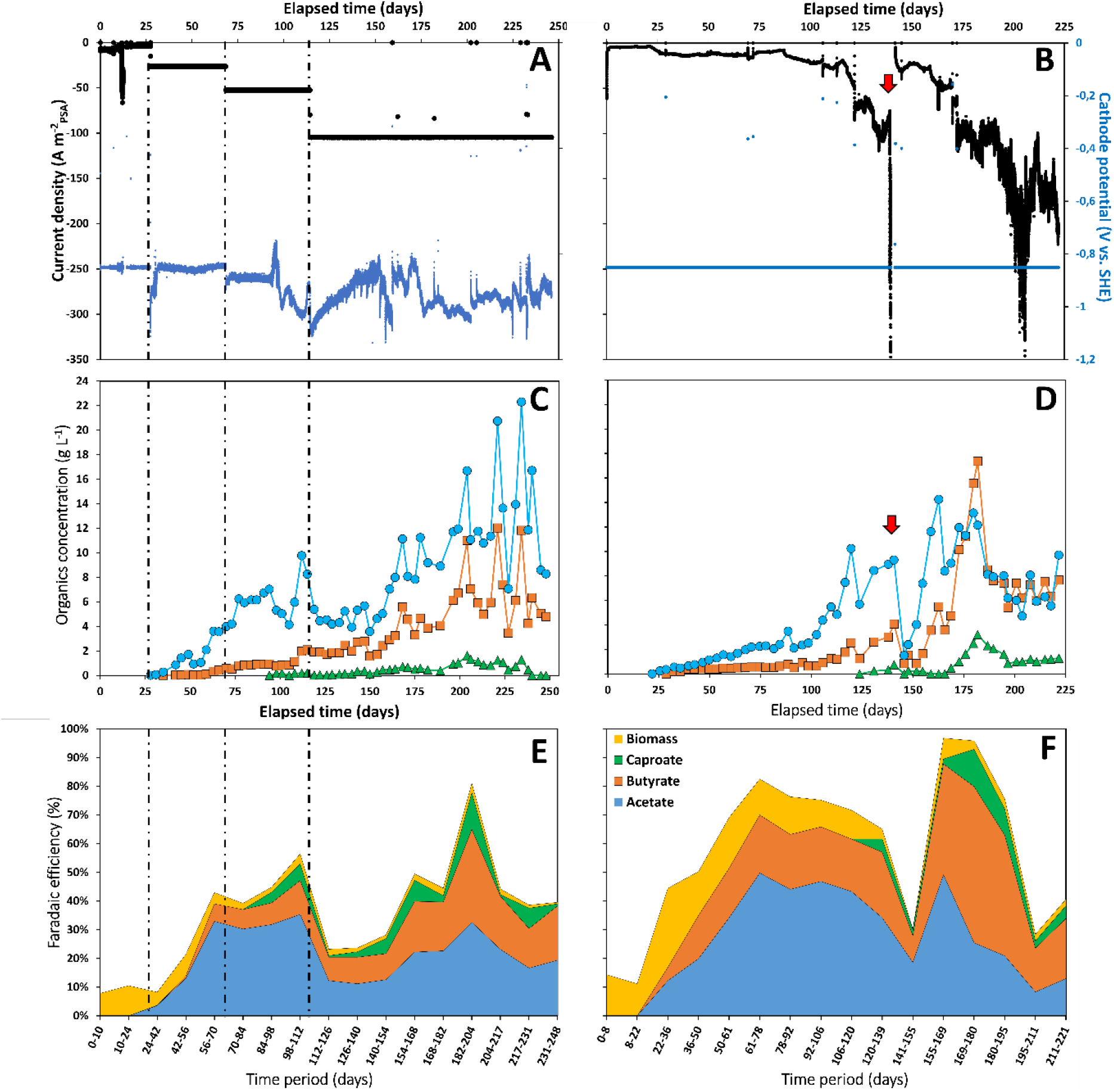
Current density and cathode potential (A-B), concentration of acetate (blue circles), butyrate (orange squares) and caproate (green triangles) (C-D), and faradaic efficiency (E-F) evolution over time in the galvanostatic (left) and potentiostatic (right) reactors. The red arrow (day 139) indicates a leakage event that emptied the cathode compartment almost entirely, which stopped the potentiostat and the liquid recirculation, and the reactor remained in this state for 3 days (long weekend).

In the galvanostatic reactor, the cathode potential consistently ranged between -0.8 V and -1.0 V throughout successive applied current steps, reaching up to -105 A m^-2^_PSA_ (projected surface area). In the potentiostatic reactor (-0.85 V vs. SHE), the cathodic current initially remained low at around -5 A m^-2^_PSA_ for the first 25 days, gradually increasing to -15 A m^-2^_PSA_ by day 35. Subsequently, the current remained relatively stable until day 85, coinciding with approximately 30% biofilm colonization of the carbon felt (**Figure 2B**). Between day 85 and day 139, the current exhibited exponential growth, reaching -100 A m^-2^_PSA_, corresponding to 50% biofilm coverage on the carbon felt. The reactor experienced a crash on day 139, inducing biofilm stress, resuspension, and rapid re-attachment, evident by a sudden increase and subsequent decrease in optical density (Figure S6). Following a lag phase, the electron uptake rate recovered from day 170 and remarkably surged further to a cathodic current of approximately -200 A m^-2^_PSA_ (-40 kA m^-3^_cathode_) at the experiment’s conclusion, with peaks reaching -300 A m^-2^_PSA_ (-60 kA m^-3^_cathode_) over a 7-day period between days 200 and 207. The prior state-of-the-art MES reactor, producing soluble organics, reported a current of - 101 A m^-2^_PSA_, equivalent to -7.8 kA m^-3^_cathode_, at the same cathode potential.^6^ The DFBR demonstrated a notable enhancement, achieving a two-fold increase in current normalized to projected surface area and a five-fold increase in volume of the electrode. Normalizing performance to electrode volume is particularly relevant when employing 3D electrodes,^33, 34^ accounting for the electrode thickness. The previous state of the art current of -101 A m^-2^_PSA_ was attained with a 1.2 cm thick carbon felt, whereas our study employed a 0.5 cm thick carbon felt. A thinner cathode is advantageous, promoting lower ohmic resistances and consequently higher energy efficiencies.^2, 11^ In the last 10 days of the experiment, a faradaic efficiency of 40% (and increasing) was achieved, corresponding to volumetric productivities of 43 kg_C2_ m^-3^_cathode_ d^-1^, 30 kg_C4_ m^-3^ _cathode_ d^-1^, and 5 kg_C6_ m^-3^_cathode_ d^-1^ (equivalent to a total C production of 37.3 kg_C_ m^-3^_cathode_ d^-1^).

### Biofilm suspension and re-attachment leads to higher carbon selectivity and faradaic efficiency into C4 and C6

Within the initial 25 days, only biomass growth occurred, representing approximately 11.3 ± 2.9% of electron recovery into biomass in both reactors, at comparable current densities. Consequently, both reactors exhibited a lag phase of approximately 25 days before measurable amounts of soluble organics were produced, a phenomenon previously observed in microbial electrosynthesis (MES).^28, 35^ Acetate (C2) was the initial product in both reactors, closely followed by n-butyrate (C4). During the initial 100 days, under galvanostatic control, C2 and C4 concentrations increased more rapidly than under potentiostatic control (7.0 vs. 2.5 g_C2_ L^-1^ and 0.9 vs. 0.6 g_C4_ L^-1^), albeit at the expense of faradaic efficiency (45% vs. 75%). A lower faradaic efficiency corresponds to reduced energy efficiency (Figure S7), indicating higher energy wastage. Despite the application of a higher current (CP) than that measured under CA, it did not result in increased growth rates or higher biofilm amounts (28% electrode colonization in both reactors at day 100, **Figure 2**). It did however lead to higher acetate production rates in the first 100 days, 26 vs 10 kg_C2_ m^-3^_cathode_ d^-1^ (Figure S4) suggesting a decoupling of growth and production metabolisms. As observed previously, once a mature biofilm was established, carboxylate production became maintenance-driven.^28^ This pattern persisted here, with acetate concentration reaching up to 10 g L^-1^ and productivity 34 kg_C2_ m^-3^_cathode_ d^-1^ when the current density increased to similar, and subsequently higher, levels under CA from day 125 to 139 (up to -100 A m^-2^). Simultaneously, butyrate concentration reached 4 g L^-1^ and productivity 18 kg_C4_ m^-3^_cathode_ d^-1^. A higher faradaic efficiency of 65% was achieved at -100 A m^-2^ under CA compared to -52 A m^-2^ under CP (50%). Caproate (C6) production commenced earlier under CP than CA (98 vs. 130 days), likely attributable to the earlier attainment of higher concentrations of C2 and C4.^6, 18^

Following the lag phase subsequent to the CA reactor crash on day 139, a notable faradaic efficiency of 90% was achieved from day 155 to 169 at a current density of approximately -47 A m^-2^. An even higher faradaic efficiency of 93% was reached at -102 A m^-2^_PSA_ (-20 kA m^-3^_cathode_) from day 169 to 180, coinciding with elevated concomitant volumetric productivities of 50 kg_C2_ m^-3^_cathode_ d^-1^, 71 kg_C4_ m^-3^_cathode_ d^-1^, and 15 kg_C6_ m^-3^_cathode_ d^-1^ (equivalent to 69 kg_C_ m^-3^_cathode_ d^-1^). This represents a remarkable 3.1-fold increase in soluble organics productivity compared to the state-of-the-art (22 kg_C_ m^-3^_cathode_ d^-1^).^6^ Notably, a high carbon selectivity of 57%C4 and 14%C6 (Figure S8) and a faradaic efficiency of 55%C4 and 13%C6 into C4 and C6, respectively, were achieved. Higher selectivity towards C4 and C6 is advantageous given their higher value compared to acetate.^4^ Lower carbon selectivity (29%_C4_ and 4%_C6_) and faradaic efficiency (23%_C4_ and 5%_C6_) into C4 and C6 were recorded before the reactor crash at the same current density of -102 A m^-2^_PSA_ from day 120 to 139. Similarly, a lower carbon selectivity (36 ± 6%C4 and 8 ± 5%C6) and faradaic efficiency (16 ± 7%C4 and 4 ± 4%C6) into C4 and C6 were achieved in the CP reactor at -105 A m^-2^, which did not experience significant biofilm disturbance. A previous study also demonstrated that rapid detachment and reattachment, leading to biofilm reorganization, significantly improved carbon selectivity and faradaic efficiency towards C4 and C6 over acetate.^18^ The mechanism responsible for this phenomenon is yet to be fully elucidated. It is noteworthy that a lower total faradaic efficiency was recorded from day 120 to 139 (62%) compared to day 169 to 180 (93%) at the same current density in the CA reactor. This discrepancy may be attributed to a lower biofilm amount (7.4 vs. 10.8 mmol_x_ cm^-3^_cathode_) and electrode coverage (43 vs. 68%) (**Figure 2B**) and/or to the biofilm reorganization.

The faradaic efficiency declined notably after the escalation of current density to -200 A m^-2^_PSA_ (-40 kA m^-3^_cathode_) from day 195 to 221. The cause of this surge in cathodic current remains unclear at this stage, but it might be attributed to the increase in electrode coverage by biofilm to over 90% during that period (**Figure 2B**). A higher biofilm amount correlates with an increased demand for electrons. On day 180, the highest concentrations ever reported in continuously-operated MES reactors of C4 (17.4 g L^-1^) and C6 (3.2 g L^-1^) were reached. Subsequently, a decline in organics concentration and production rates was observed. A plausible explanation could be the inhibition of the biofilm by C4 and C6 acids, known for their toxicity, as previously modeled in MES,^13^ though further investigation is required for confirmation. Nonetheless, a 2-fold higher volumetric productivity compared to the previous state-of-the-art was maintained from day 195 until the experiment’s conclusion. Additionally, carbon selectivity remained favorable, with 57% and 11% directed into C4 and C6, respectively.

### Potentiostatic control results in higher faradaic and energy efficiencies

Under galvanostatic control, an increment in cathodic current density from -26 to -53 or -105 A m^-2^ did not significantly elevate the organics concentration until day 150 (**Figure 4C**). This observation could be attributed to the fact that by day 150, only half of the electrode was colonized (**Figure 2B**). The limited biofilm coverage hampers the microbial uptake of electrons (and CO_2_). This is evident in the relatively low faradaic efficiency during this period. Subsequently, from day 150 until the end of the experiment, concentrations and production rates gradually increased as biofilm coverage expanded. High acetate and butyrate concentrations of 22.3 g L^-1^ and 12 g L^-1^ were attained in this reactor. Simultaneous volumetric productivities of 64 kg_C2_ m^-3^_cathode_ d^-1^, 42 kg_C4_ m^-3^_cathode_ d^-1^, and 17 kg_C6_ m^-3^_cathode_ d^-1^ (equivalent to 60 kg_C_ m^-3^_cathode_ d^-1^) were achieved (Figure S4), marking a 2.7-fold increase compared to the previous state-of-the-art and similar to the CA reactor. Relative to the CA reactor, lower faradaic and energy efficiencies were observed under galvanostatic control, despite similar current density and biofilm amount/coverage. The reasons for these differences warrant further investigation. In the CA reactor, energy efficiencies averaged 34 ± 17% at -102 A m^-2^, with a peak at 64% (Figure S7). It is important to note that optimizing the energy efficiency of the directed-flow-through bioelectrochemical reactor (DFBR) was beyond the scope of this work; for instance, a thick membrane was employed, leading to high ohmic overpotentials. Overall, galvanostatic control did not accelerate biofilm growth or result in higher organics productivity and faradaic efficiency in this study.

### Biomass-specific production rates maintained over a long period of time

Employing a recently established methodology,^28^ biomass-specific production rates (*q*_p_) can now be determined over time in MES (**Figure 5**). Biomass-specific production rate represents a microbial kinetic parameter that normalizes the production rate to the quantity of biomass within the reactor at a specific moment. Utilizing biomass-specific rates, one can more effectively assess the specific performance of the biocatalysts in MES, facilitating meaningful comparisons with other technologies.

In both reactors, *q*_p_ exhibited an initial increase before stabilizing at comparable values of 0.19 ± 0.06 mol_C_ mol_x_^-1^ d^-1^ until the conclusion of the experiments. A preceding study, which reported *q*_p_ in microbial electrosynthesis, documented biomass-specific production rates of C2-C6 carboxylates around 0.25 ± 0.05 mol_C_ mol_x-_ ^-1^ d^-1^ in the initial 50 days, followed by a gradual decline to 0.05 ± 0.03 mol_C_ mol_x-_^-1^ d^-1^ by day 200.^28^ It was suggested that this rate, like the specific growth rate, declined as the biofilm quantity increased and matured to fully colonize the electrode. The aforementioned study utilized non-optimized flow-through reactors with the same electrode material and inoculum. In contrast, our findings demonstrate that the novel directed-flow-through bioelectrochemical reactor concept allows for the preservation of biomass-specific production rates over an extended period, even with biofilm apparent density three times higher and elevated product concentrations compared to the study by Winkelhorst et al.^28^ These results can be attributed to improved mass transport of CO_2_, nutrients, H_2_, and/or products throughout the entire cathode, facilitated by the specific reactor architecture and fluid dynamics.

### Volumetric productivity in MES now comparable to syngas fermentation

Syngas fermentation serves as a pertinent benchmark for assessing microbial electrosynthesis. Syngas fermentation, employing acetogens to convert gas mixtures of H_2_, CO, and CO_2_ into a mixture of acetate and ethanol, has been successfully scaled up to an industrial level, exemplified by the LanzaTech process.^7^ When comparing these technologies, three key performance indicators— biomass-specific production rates (*q*_p_), the amount of biomass per reactor volume, and the product of both i.e. the volumetric productivity—should be considered. A comparison was drawn using data from two lab-scale syngas fermentation studies reporting the highest performance to the best of our knowledge,^8, 9^ and the industrial-scale LanzaTech process (**Figure 5B**).^7^ The achieved biomass-specific production rates in MES were notably lower than those reported in syngas fermentation, reaching up to 20 mol_C_ mol_x_ ^-1^ d^-1^. This observation presents an opportunity for MES, indicating the potential for higher microbial rates. Conversely, the amount of microbial biomass per unit of reactor volume in MES (390 g_x_ L^-1^ _cathode_) significantly surpassed that in syngas fermentation (2.5 g_x_ L^-1^). MES relies on a dense biofilm, while the highest syngas fermentation performances were attained with microorganisms in suspension. Consequently, our MES process demonstrated a comparable volumetric productivity of approximately 0.2 g_C_ L^-1^ h^-1^ to lab-scale syngas fermentation and was five times lower than the LanzaTech process. It is important to note that volumetric productivity calculations in MES here utilized the total volume of catholyte. Normalizing to the volume of catholyte in the cathode chamber yielded productivity values 14 times higher, surpassing the LanzaTech process by 2.5 times. Optimization of the catholyte to electrode volume ratio requires further investigation. Additionally, considering market prices, it is noteworthy that C4 and C6 carboxylic acids, produced in MES, have higher prices than ethanol, although their market volumes are lower.^4^ This study represents a milestone in developing MES as a competitive technology, marking a promising outcome that warrants further scaling-up of MES technology.

Furthermore, microbial rates in microbial electrosynthesis are within the same order of magnitude as rates achieved in gas fermentation converting H_2_ + CO_2_ to carboxylates.^36, 37^ In a study by Zhang et al.,^36^ biofilms were cultivated on hollow-fibre membranes to convert an H_2_/CO_2_ mixture, yielding volumetric production rates considerably lower at 0.003 g_C_ L^-1^ h^-1^ compared to the rates reported in our study. A systematic comparison of these two technologies is essential to delineate the advantages and disadvantages of each approach, especially in the context of scale-up considerations. Notably, Roghair et al. reported higher biomass-specific rates of 10 mol_C_ mol_x_ ^-1^ d^-1^ in their fermentation process, converting acetate and ethanol to a mixture of C2-C6 carboxylates, accompanied by a volumetric productivity of 2.7 g_C_ L^-1^ h^-1^.^38^ Such comparative analyses can contribute valuable insights into optimizing and advancing these microbial processes.

It is important to note that the calculated biomass-specific production rates (*q*_p_) in MES presented in this study are underestimated, as they do not consider the accumulation of dead biomass within the biofilm. Winkelhorst et al.^28^ demonstrated that after 200 days, up to 50% of the microbes within the biofilms were found to be dead. Consequently, the estimated *q*_p_ values were approximately half of the actual values, emphasizing the need to account for dead biomass to obtain a more accurate assessment of microbial kinetics in MES.

### *Clostridium luticellarii* and *Eubacterium limosum* are dominant species

To gain a deeper understanding of the notable performance distinctions between potentiostatically (CA) operated bioreactors and their galvanostatically (CP) controlled counterparts, we employed high-throughput 16S rRNA gene sequencing to investigate microbial community structures. Following an experimental period exceeding 220 days, microbial samples were collected from cathode-attached cells at three locations along the serpentine flow channel (Figure 6A). Each sample exhibited a dominant amplicon sequence variant (ASV) representation of more than 65%, encompassing no more than seven ASVs. Across the six samples, only 13 dominant species were identified (Figure 6B). Four species—*Eubacterium limosum, E. callanderi, Fermentimonas caenicola*, and *Clostridium luticellari*—were consistently present in all samples, regardless of the operational mode.

**Figure 5.**
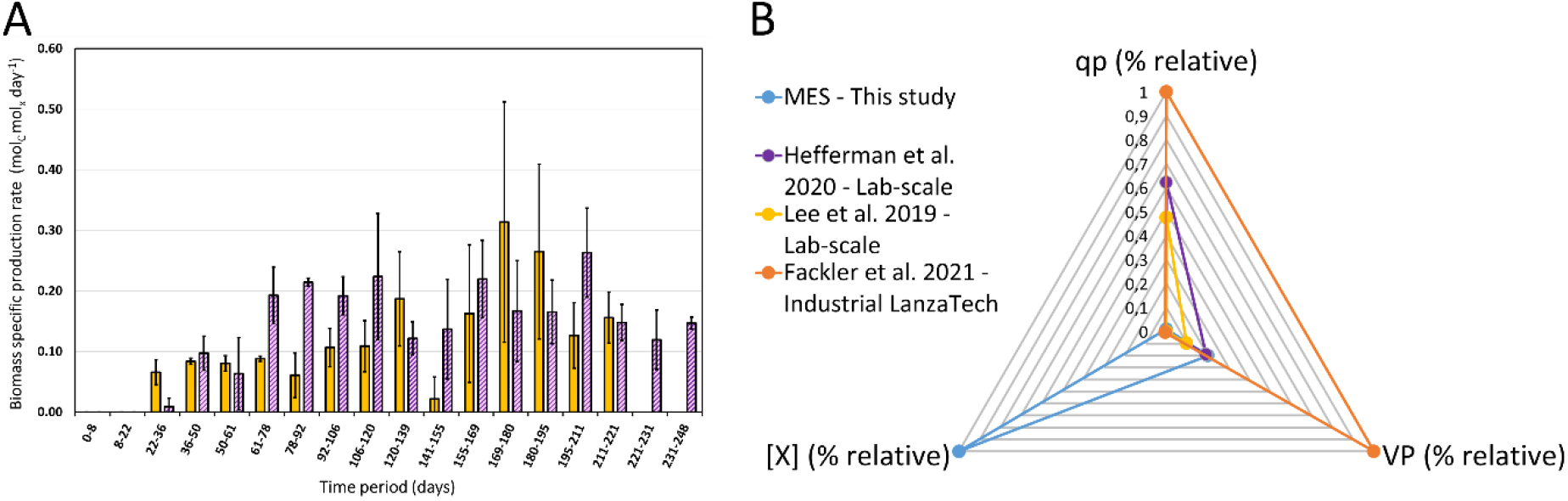
A) Biomass-specific production rates measured over time in the galvanostatic (purple bars) and potentiostatic (yellow bars) reactors. B) Comparison of three key performance indicators between our MES reactor and two of the best-performing lab-scale syngas fermentation reported in literature^8, 9^ as well as the industrial LanzaTech^7^ syngas fermentation process. Instead of absolute values for each KPIs, relative percentages calculated from the max value are represented. Biomass specific production rates q_p_ vary from 0,2 to 20 mol_C_ mol_x-1_ d^-1^, biomass concentrations [X] from 0.5 to 390 g_x_ L^-1^, and volumetric productivities (VP) from 0.1 to 1 g_C_ L^-1^ h^-1^. VP is here normalized to the total medium volume, and not to the cathode volume in MES. Refer to Table S1 in the supplementary file to see the values extracted from literature and used here.

**Figure 6.**
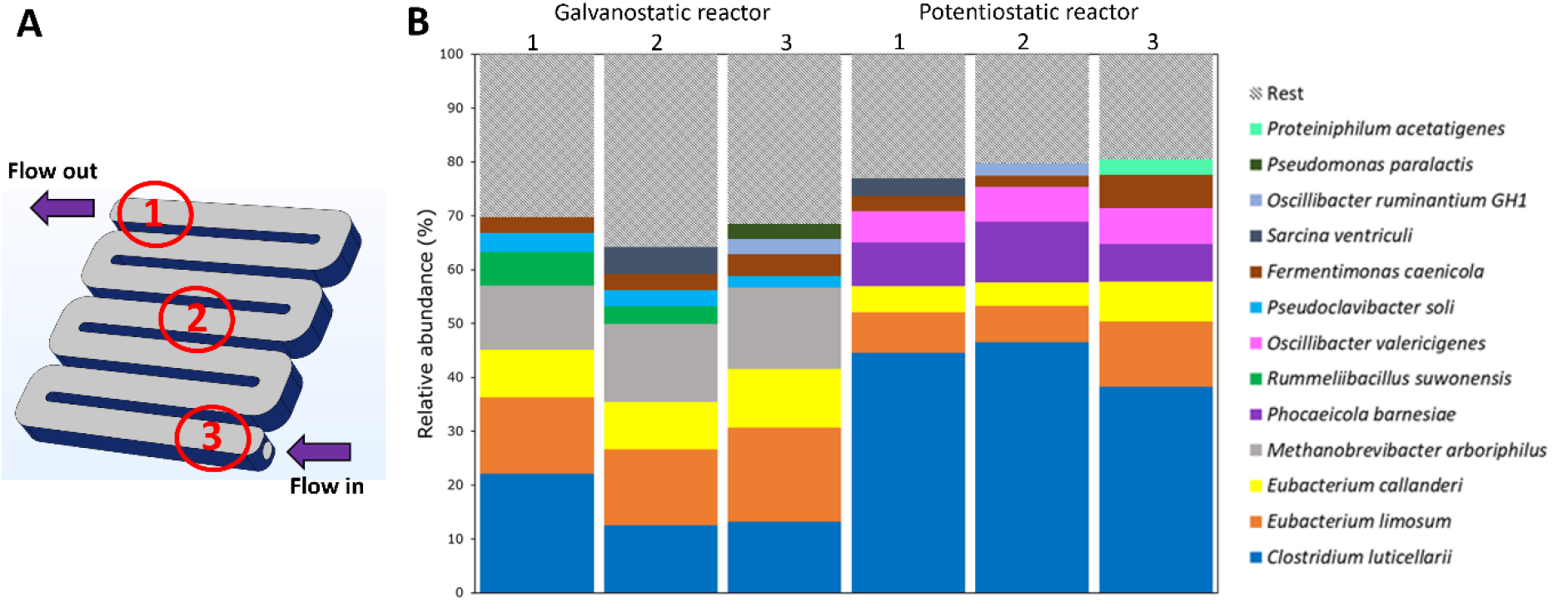
A. Sampling location along the serpentine electrode. B. Relative abundance of 16S rRNA of biofilm samples taken from three locations in both the galvanostatic and potentiostatic reactors, at the end of the experiment.

Among the remaining nine species, eight were specific to a single process. The galvanostatic reactor consistently featured ASVs belonging to the *Methanobrevibacter* and *Pseudoclavibacter* genera. In contrast, microbial communities from the potentiostatic reactor consistently shared six species, including *Oscillibacter* and *Phocaeicola* ASVs, in addition to the four species systematically found. Strikingly, none of these 13 ASVs have been associated to MES microbial communities before.^39-41^

Three of the four systematically identified dominant species—*E. limosum, E. callanderi*, and *Clostridium luticellari* belong to the Clostridia class. These are known acetogens typically found in microbial communities associated with syngas fermentation. In this process, they grow autotrophically by utilizing a mixture of H_2_, CO and CO_2_ as carbon and energy sources, converting them into the central intermediate acetyl-CoA which is converted to acetate by most acetogens.^42^ Interestingly, *Eubacterium limosum* and *Clostridium luticellari* were also found to perform chain elongation and produce C4 and C6 carboxylic acids,^42, 43^ a characteristic dependent on environmental conditions. The presence of the three anaerobic species *Phocaeicola barnesiae*,^44^ *Oscilibacter valericigenes and Fermentimonas caenicola* remains unclear. Only *Oscilibacter valericigenes* (*Clostridia* class) was linked to C5 carboxylic acid biosynthesis.^45^ The presence of these ancillary species could be necessary to meet specific nutritional requirements and contribute to the stable establishment of the biofilm.

On basis of these initial taxonomic findings, it is inferred that the serpentine design, which establishes a flow pattern in the forced-flow-through reactor, enabling an extended residence time for nutrients, CO_2_, H_2_, and products within the 3D-electrode, did not yield a pronounced gradient in community composition along the flow channel in each reactor. The microbial composition was found to be comparable at all three locations. It is essential to underscore that relative abundance does not necessarily align with activity levels. To understand the microbial function and physiology of these highly active biofilms, further analyses involving metagenomics, metatranscriptomics, and metaproteomics are imperative.

The source of the biofilm inoculum plays a pivotal role in shaping the composition of the ultimate microbial community, and the considerable diversity in inoculum origins documented in scientific literature poses challenges for meaningful comparisons. Nonetheless, our results conclusively demonstrate that novel microbial communities capable of driving highly efficient processes can be assembled in MES bioreactors.

### MES: a competitive technology for efficient CO_2_ valorization

We established that forcing an electron flow through the application of current does not accelerate the growth of biofilms. This investigation underscores that the primary constraint on the applicability of microbial electrosynthesis lies in the rate at which a large electrode can be colonized. Once colonization occurs, we demonstrated the attainment of high current densities, productivities, and faradaic efficiencies. The designed directed-flow-through bioelectrochemical reactor (DFBR) showcased in this work enables the achievement of elevated biofilm concentrations per electrode volume, sustained microbial activity over a period exceeding 225 days, and notable volumetric productivity, now comparable to rates observed in syngas fermentation. This study emphasizes the feasibility of enhancing microbe-specific rates to further increase volumetric productivity and product selectivity. Our findings represent a milestone in developing MES as a competitive technology for efficient CO_2_ valorization.

## Experimental Procedures

### Resource availability

#### Lead Contact

Further information and requests for resources should be directed to and will be fulfilled by the Lead Contact, Ludovic Jourdin.

#### Materials Availability

This study did not generate new unique materials.

#### Data and Code Availability

The data underlying the publication are publicly available on 4TU.ResearchData, DOI 10.4121/28d4cf06-85e2-4c63-a632-179ede79251f, in line with the TU Delft Research Data Framework Policy. Metagenomic V3-V4 16S rRNA gene Amplicon DNA-sequencing data of cathode microbiome were deposited at NCBI (https://www.ncbi.nlm.nih.gov/) under BioProject accession number PRJNA1071877.

### MES Reactor Setup

Two bioelectrochemical reactors were assembled. Each reactor comprised two identical cathodic and anodic compartments, as well as two supporting plates used to press and close the reactor cell. A scheme of the reactor setup and a picture of the assembled reactor can be seen in Figure 1. Exact reactor dimensions are also given (Figure S9). A biocompatible resin (BioMed Clear Resin V1, Formlabs) and the Form 2 printer (Formlabs) were used to 3D-print all the plates that formed the reactor.

The anode used in this work was a Pt/IrO_2_ coated titanium plate (Magneto Special Anodes, Schiedam, The Netherlands). Unmodified carbon felt was used as the cathode electrode material (CTG Carbon GmbH, Germany). Before being used, all felt pieces underwent a cleaning step with 1 mol L^-1^ HCl, 1 mol L^-1^ NaOH, and an UV/ozone treatment as described in Winkelhorst et al.^28^ Once in the reactor, the total volume of carbon felt was 9.5 cm^3^, with a thickness of 0.5 cm and a total projected surface area of 19 cm^2^. In one of the reactors, a titanium wire (Salomon’s Metalen, The Netherlands) was placed between two layers of tightly pressed carbon felt (0.25 cm thick each) and used as current collector. The other reactor used an iso-molded graphite plate (GP) (3.2 mm thick, Fuel Cell Store, USA) as current collector, placed parallel to the cathodic plate. Conductive coating (Graphite Conductive Adhesive, Electron Microscopy Sciences, USA) was applied at the surface between current collector and carbon felt in order to enhance the electric connection between both materials.^28^ A cation exchange membrane (CEM) (CMI-7000s, Membrane International Inc.) was used to separate the cathodic and anodic compartment.

A pH probe (QP108X, ProSense) was installed in the catholyte recirculation loop outside the reactor and a pH controller (AQUIS touch S, Jumo) was used to control the pH of the catholyte at pH 5.8. A bubble column was also installed in the recirculation loop, and used to sparge a CO_2_-N_2_ mixture into the catholyte. The total volume of the catholyte in the setup was 135 mL. This volume includes the cathodic chamber, recirculation bottle, and all tubing comprising the recirculation loop.

### MES Reactor Operation

The catholyte medium composition was based on the one described in Winkelhorst et al.,^28^ and contained 0.2 g L^-1^ NH_4_Cl, 0.015 g L^-1^ CaCl_2_·2H_2_O, 0.04 g L^-1^ MgCl_2_·6H_2_O, 8.1 g L^-1^ KH_2_PO_4_, 0.9 g L^-1^ Na_2_HPO_4_, 1 mL L^-1^ of a trace elements solution, and 4.5 g L^-1^ 2-bromoethanesulfonic acid as methanogenic activity inhibitor. The trace elements solution contained 1.5 g L^-1^ FeCl_3_·6H_2_O, 0.15 g L^-1^ H_3_BO_3_, 0.03 g L^-1^ CuSO_4_·5H_2_O, 0.18 g L^-1^ KI, 0.12 g L^-1^ MnCl_2_·4H_2_O, 0.06 g L^-1^ Na_2_MoO_4_·2H_2_O, 0.12 g L^-1^ ZnSO_4_·7H_2_O, 0.15 g L^-1^ CoCl_2_·6H_2_O, 0.023 g L^-1^ NiCl_2_·6H_2_O, and 10 g L^-1^ EDTA. To avoid possible limitations caused by nutrients depletion, a second catholyte medium composition was used from day 18 containing 0.6 g L^-1^ NH_4_Cl, 0.045 g L^-1^ CaCl_2_·2H_2_O, 0.12 g L^-1^ MgCl_2_·6H_2_O, 8.1 g L^-1^ KH_2_PO_4_, 0.9 g L^-1^ Na_2_HPO_4_, 3 mL L^-1^ of a trace elements solution, and 13.5 g L^-1^ 2-bromoethanesulfonic acid. The anolyte composition was identical to the catholyte medium used at that specific moment, without the trace elements solution and the methanogenic inhibitor. The pH of the anolyte was also decreased to 2 with phosphoric acid to facilitate proton crossing over the membrane.

The reactors were operated in continuous mode with a hydraulic retention time (HRT) of 4 days. Both catholyte and anolyte were continuously circulated between the reactor cell and the recirculation bottles at a flow rate of 1.8 L h^-1^. Dissolved CO_2_ was supplied by continuously sparging a gas mixture of CO_2_:N_2_ 50:50 at a rate of 0.1 L min^-1^ in the cathodic bubble column. The volumetric CO_2_ mass transfer coefficient (k_L_a) of the bubble column was determined with the dynamic gassing-out method and found to be 70 h^-1^ (data not shown). The entire setup was placed inside a cabinet, keeping the reactors in the dark to avoid any phototrophic growth. Temperature was kept at 31 °C.

The reactors were connected in a three-electrode configuration to a multichannel potentiostat (BioLogic) to control either the cathode potential (chronoamperometry, CA) or the applied current (chronopotentiometry, CP). A 3M Ag/AgCl reference electrode (QM710X, ProSense) was installed in both reactors. The CP reactor was first controlled in potentiostatic mode at -0.85 V vs. SHE (standard hydrogen electrode) from day 0 to 28, and then switched to galvanostatic mode for the rest of the experiment (day 248). From day 28 to 68 a current of -26 A m_2_ ^-1^_psa_ was applied, from day 68 to 114 it was increased to -53 A m_2_^-1^_psa_, and from day 114 to the end of the experiment -105 A m_2_^-1^_psa_ were applied. The CA reactor was controlled at -0.85 V vs. SHE throughout the course of the experiment (221 days).

Each reactor was inoculated on day 0 with 1.2 g COD_x_ L^-1^. The inoculum was taken from running laboratory MES reactors producing acetate, *n*-butyrate, and *n*-caproate from CO_2_.^28^

### Analytical Methods

Twice a week, liquid samples were taken from each reactor. Concentration of C2 to C6 carboxylic acids and alcohols were analyzed by gas chromatography (Thermofisher, USA) using a Stabil-wax™ column with a length of 25 m and internal diameter of 0.2 μm. Colum temperature was kept at 50 °C for 7 min, increased to 180 °C during 8 min and kept at that temperature for 9 minutes. Helium was used as carrier gas at a flow rate of 1 mL min^-1^ and the ionization detector was kept at 250 °C. Production of organics and faradaic efficiency were calculated as described by Raes et al.^46^ Biomass-specific growth rates and production rates were calculated as described by Winkelhorst et al.^28^ Here, carbon selectivity represents the fraction of carbon going into a specific product over the total amount of carbon assimilated in all identified products, i.e. acetate, butyrate, caproate, and biomass.

Samples were filtered with a 0.2 μm microporous filter and their total nitrogen content was analysed using a TOC analyser coupled with a TN unit (TOC-L Series Total Organic Carbon Analyzers, Shimadzu, Japan). Oven temperature was kept at 720 °C. The optical density of the unfiltered samples was measured at 600 nm with a UV-VIS spectrophotometer (UV-1800 series, Shimadzu, Japan) to account for planktonic cells. To study microbial activity, the method described by Winkelhorst et al.^28^ was utilized to estimate microbial growth in both biofilm and suspension separately as well as biomass concentration and biomass-specific productivity.

### Scanning Electron Microscopy

Carbon felt pieces were carefully cut at different places along the length of the flow channel (Figure 6). Samples were immediately fixed with 2.5% glutaraldehyde in PBS buffer for 24 h at 4 °C, rinsed with PBS buffer and dehydrated with a graded series of ethanol.

### DNA extraction and 16S sequence analysis

To extract DNA from the biofilm both on and within the 3D porous cathode, the samples were ground with a pestle in a mortar, periodically incorporating liquid nitrogen to prevent DNA degradation. The resulting powder comprised a mixture of biomass and carbon felt fibers. Subsequently, DNA extraction was carried out using a Qiagen DNeasy PowerBiofilm Kit (Qiagen, Hilden, Germany), employing a detergent lysis method in conjunction with bead beating using an MPTM FastPrep-24 homogenizer (MP Biomedicals, Irvine, CA), following the manufacturer’s instructions. The purified DNA underwent quality assessment by measuring the A260/280 and A260/230 ratios (Nanodrop 2000 spectrophotometer (ThermoFischer Scientific, Waltham, MA) and quantification using a Qubit broad-range assay (Qubit 2.0 Fluorometer and Qubit dsDNA BR Assay Kit (Thermo Fischer Scientific)).

### Microbial Community Analysis

Microbial community analysis was conducted through 16S rRNA sequencing. For this purpose, extracted DNA samples were sent to Novogene (UK) (Cambridge, United Kingdom). The 16S rRNA amplicon was sequenced using barcoded primers 341F (CCTAYGGGRBGCASCAG) and 806R (GGACTACNNGGGTATCTAAT) to amplify regions V3 + V4 of both bacterial and archaeal microorganisms. All polymerase chain reactions (PCR) were carried out in 30 μL reaction volumes, comprising 15 μL of Phusion® High-Fidelity PCR Master Mix (New England Biolabs, Ipswich, MA), 0.2 μM of forward and reverse primers, and approximately 10 ng of template DNA. Thermal cycling involved initial denaturation at 98 °C for 10 s, annealing at 50 °C for 30 s, elongation at 72 °C for 30 s, and a final extension at 72 °C for 5 min. PCR products underwent purification through gel electrophoresis using a 2% agarose gel and subsequent extraction using a Qiagen Gel Extraction Kit. Sequencing libraries were generated using the NEBNext® UltraTM DNA Library Prep Kit for Illumina (Illumina, San Diego, CA), following the manufacturer’s recommendations, with index codes added. The library quality was evaluated using a Qubit 2.0 Fluorometer (Thermo Scientific) and Agilent Bioanalyzer 2100 system (Agilent, Santa Clara, CA), and the sequencing was performed on an Illumina HiSeq platform to generate 250 bp paired-end reads. Data analysis was performed as described in Supplemental information.

## Supplemental information description

**Figure S1**. Biofilm concentration measured over time in the galvanostatic (orange triangles) and potentiostatic (blue circles) reactors, compared with the state-of-the-art from Winkelhorst et al. 2023 (green diamonds).

**Figure S2**. Relative abundance of biomass in biofilm and in suspension in the galvanostatic (A) and potentiostatic (B) reactors.

**Figure S3**. Photograph of the biofilm on the side of the carbon felt facing the current collector (A) and throughout its thickness (B). SEM images of the biofilm grown on the carbon felt electrode of the potentiostatic (C-D) and galvanostatic (E-F) reactors, at the end of the experiment.

**Figure S4**. Production rates, normalized to electrode volume (A-B) and total catholyte volume (C-D), of acetate (blue circles), butyrate (orange squares) and caproate (green triangles) measured over time in the galvanostatic (A, C) and potentiostatic (B, D) reactors.

**Figure S5**. Current profile in kA m^-3^_cathode_ in the galvanostatic (A) and potentiostatic (B) reactors.

**Figure S6**. Soluble nitrogen concentration (A-B), optical density and normalized overpressure (C-D) measured over time in the galvanostatic (A & C) and potentiostatic (B & D) reactors.

**Figure S7**. Energy efficiency over time in the galvanostatic (A) and potentiostatic (B) reactors.

**Figure S8**. Carbon selectivity over time in the galvanostatic (A) and potentiostatic (B) reactors.

**Figure S9**. Reactor scheme with dimensions (left and middle) and photo of the reactor (right).

**Table S1**. Literature data used to make Figure 5B.

**Sequencing data analysis**.

## Acknowledgements

This activity was co-financed by Shell and a PPP-allowance from Top Consortia for Knowledge and Innovation (TKI’s) of the Dutch Ministry of Economic Affairs and Climate in the context of the TU Delft e-Refinery Institute. RS, JMD, and LJ acknowledge the co-financing by DSM-Firmenich and a PPP-allowance from Top Consortia for Knowledge and Innovation (TKI’s) of the Dutch Ministry of Economic Affairs and Climate.

## Author contributions

OCP, MW, and RKA operated the MES reactors. OCP and LJ designed the study, designed the novel reactor, analyzed and interpreted the data, and drafted the manuscript. RS and JMD analyzed the microbial communities and drafted the manuscript. KM 3D-printed the reactors. LJ, JMD, and AJJS acquired the funding. All authors contributed to manuscript revision, conception, and read and approved the submitted version.

## Declaration of interest

LJ and OCP have a patent pending related to this work (application number NL2032221).

## Table titles and legends

Not applicable.

## References

1. Rabaey, K. and R.A. Rozendal, Microbial electrosynthesis — revisiting the electrical route for microbial production. Nat Rev Micro, 2010. 8(10): p. 706–716.

2. Jourdin, L. and T. Burdyny, Microbial Electrosynthesis: Where Do We Go from Here? Trends in Biotechnology, 2020.

3. Flexer, V. and L. Jourdin, Purposely Designed Hierarchical Porous Electrodes for High Rate Microbial Electrosynthesis of Acetate from Carbon Dioxide. Accounts of Chemical Research, 2020. 53(2): p. 311–321.

4. Jourdin, L., J. Sousa, N.v. Stralen, and D.P.B.T.B. Strik, Techno-economic assessment of microbial electrosynthesis from CO2 and/or organics: An interdisciplinary roadmap towards future research and application. Applied Energy, 2020. 279: p. 115775.

5. Vassilev, I., P.A. Hernandez, P. Batlle-Vilanova, S. Freguia, J.O. Krömer, J. Keller, P. Ledezma, and B. Virdis, Microbial Electrosynthesis of Isobutyric, Butyric, Caproic Acids, and Corresponding Alcohols from Carbon Dioxide. ACS Sustainable Chemistry & Engineering, 2018. 6(7): p. 8485–8493.

6. Jourdin, L., S. Raes, C. Buisman, and D. Strik, Critical biofilm growth throughout unmodified carbon felts allows continuous bioelectrochemical chain elongation from CO2 up to caproate at high current density. Frontiers in Energy Research, 2018. 6: p. 7.

7. Fackler, N., B.D. Heijstra, B.J. Rasor, H. Brown, J. Martin, Z. Ni, K.M. Shebek, R.R. Rosin, S.D. Simpson, K.E. Tyo, et al., Stepping on the Gas to a Circular Economy: Accelerating Development of Carbon-Negative Chemical Production from Gas Fermentation. Annual Review of Chemical and Biomolecular Engineering, 2021. 12(1): p. 439–470.

8. Heffernan, J.K., K. Valgepea, R. de Souza Pinto Lemgruber, I. Casini, M. Plan, R. Tappel, S.D. Simpson, M. Köpke, L.K. Nielsen, and E. Marcellin, Enhancing CO2-Valorization Using Clostridium autoethanogenum for Sustainable Fuel and Chemicals Production. Frontiers in Bioengineering and Biotechnology, 2020. 8.

9. Lee, J., J.W. Lee, C.G. Chae, S.J. Kwon, Y.J. Kim, J.-H. Lee, and H.S. Lee, Domestication of the novel alcohologenic acetogen Clostridium sp. AWRP: from isolation to characterization for syngas fermentation. Biotechnology for Biofuels, 2019. 12(1): p. 228.

10. Nevin, K.P., T.L. Woodard, A.E. Franks, Z.M. Summers, and D.R. Lovley, Microbial electrosynthesis: Feeding microbes electricity to convert carbon dioxide and water to multicarbon extracellular organic compounds. mBio, 2010. 1(2): p. 1–4.

11. Prévoteau, A., J.M. Carvajal-Arroyo, R. Ganigué, and K. Rabaey, Microbial electrosynthesis from CO2: forever a promise? Current Opinion in Biotechnology, 2020. 62: p. 48–57.

12. Vassilev, I., P. Dessì, S. Puig, and M. Kokko, Cathodic biofilms – A prerequisite for microbial electrosynthesis. Bioresource Technology, 2022. 348: p. 126788.

13. Cabau-Peinado, O., A.J.J. Straathof, and L. Jourdin, A General Model for Biofilm-Driven Microbial Electrosynthesis of Carboxylates From CO2. Frontiers in Microbiology, 2021. 12(1405).

14. Kühl, M. and B. Jørgensen Bo, Microsensor Measurements of Sulfate Reduction and Sulfide Oxidation in Compact Microbial Communities of Aerobic Biofilms. Applied and Environmental Microbiology, 1992. 58(4): p. 1164–1174.

15. Picioreanu, C., I.M. Head, K.P. Katuri, M.C.M. van Loosdrecht, and K. Scott, A computational model for biofilm-based microbial fuel cells. Water Research, 2007. 41(13): p. 2921–2940.

16. Picioreanu, C., M.C.M. van Loosdrecht, and J.J. Heijnen, A theoretical study on the effect of surface roughness on mass transport and transformation in biofilms. Biotechnology and Bioengineering, 2000. 68(4): p. 355–369.

17. Alqahtani, M.F., K.P. Katuri, S. Bajracharya, Y. Yu, Z. Lai, and P.E. Saikaly, Porous Hollow Fiber Nickel Electrodes for Effective Supply and Reduction of Carbon Dioxide to Methane through Microbial Electrosynthesis. Advanced Functional Materials, 2018. 28(43): p. 1804860.

18. Jourdin, L., M. Winkelhorst, B. Rawls, C.J.N. Buisman, and D.P.B.T.B. Strik, Enhanced selectivity to butyrate and caproate above acetate in continuous bioelectrochemical chain elongation from CO2: Steering with CO2 loading rate and hydraulic retention time. Bioresource Technology Reports, 2019. 7: p. 100284.

19. Krieg, T., J. Madjarov, L.F.M. Rosa, F. Enzmann, F. Harnisch, D. Holtmann, and K. Rabaey, Reactors for Microbial Electrobiotechnology. 2018, Springer Berlin Heidelberg: Berlin, Heidelberg. p. 1–41.

20. Cui, K., K. Guo, J.M. Carvajal-Arroyo, J. Arends, and K. Rabaey, An electrolytic bubble column with an external hollow fiber membrane gas–liquid contactor for effective microbial electrosynthesis of acetate from CO2. Chemical Engineering Journal, 2023. 471: p. 144296.

21. Rosa, L.F.M., S. Hunger, C. Gimkiewicz, A. Zehnsdorf, and F. Harnisch, Paving the way for bioelectrotechnology: Integrating electrochemistry into bioreactors. Engineering in Life Sciences, 2017. 17(1): p. 77–85.

22. Rosa, L.F.M., S. Hunger, T. Zschernitz, B. Strehlitz, and F. Harnisch, Integrating Electrochemistry Into Bioreactors: Effect of the Upgrade Kit on Mass Transfer, Mixing Time and Sterilizability. Frontiers in Energy Research, 2019. 7.

23. Enzmann, F., F. Mayer, M. Stöckl, K.-M. Mangold, R. Hommel, and D. Holtmann, Transferring bioelectrochemical processes from H-cells to a scalable bubble column reactor. Chemical Engineering Science, 2019. 193: p. 133–143.

24. Batlle-Vilanova, P., R. Ganigué, S. Ramió-Pujol, L. Bañeras, G. Jiménez, M. Hidalgo, M.D. Balaguer, J. Colprim, and S. Puig, Microbial electrosynthesis of butyrate from carbon dioxide: Production and extraction. Bioelectrochemistry, 2017. 117: p. 57–64.

25. Romans-Casas, M., R. Blasco-Gómez, J. Colprim, M.D. Balaguer, and S. Puig, Bio-electro CO2 recycling platform based on two separated steps. Journal of Environmental Chemical Engineering, 2021. 9(5): p. 105909.

26. Roy, M., M. Saich, and S.A. Patil, Scalability of the Microbial Electro-acetogenesis Process for Biogas Upgradation: Performance and Technoeconomic Assessment of a Liter-Scale System. Energy & Fuels, 2023. 37(20): p. 15822–15831.

27. de Smit, S.M., J.J.H. Langedijk, J.H. Bitter, and D. Pbtb Strik, Alternating direction of catholyte forced flow-through 3D-electrodes improves start-up time in microbial electrosynthesis at applied high current density. Chemical Engineering Journal, 2023: p. 142599.

28. Winkelhorst, M., O. Cabau-Peinado, A.J.J. Straathof, and L. Jourdin, Biomass-specific rates as key performance indicators: A nitrogen balancing method for biofilm-based electrochemical conversion. Frontiers in Bioengineering and Biotechnology, 2023. 11.

29. Groher, A. and D. Weuster-Botz, General medium for the autotrophic cultivation of acetogens. Bioprocess and Biosystems Engineering, 2016. 39(10): p. 1645–1650.

30. Candry, P., T. Van Daele, K. Denis, Y. Amerlinck, S.J. Andersen, R. Ganigué, J.B.A. Arends, I. Nopens, and K. Rabaey, A novel high-throughput method for kinetic characterisation of anaerobic bioproduction strains, applied to Clostridium kluyveri. Scientific Reports, 2018. 8(1): p. 9724.

31. Arends, J.B.A., S.A. Patil, H. Roume, and K. Rabaey, Continuous long-term electricity-driven bioproduction of carboxylates and isopropanol from CO2 with a mixed microbial community. Journal of CO2 Utilization, 2017. 20: p. 141–149.

32. Bajracharya, S., K. Vanbroekhoven, C. Buisman, D. Strik, and P. Deepak, Bioelectrochemical Conversion of CO2 to Chemicals: CO2 as Next Generation Feedstock for the Electricity-driven Bioproduction in Batch and Continuous mode. Faraday Discussions, 2017. 202: p. 433–449.

33. Jourdin, L. and D.P.B.T.B. Strik, Electrodes for Cathodic Microbial Electrosynthesis Processes: Key-Developments and Criteria for Effective Research & Implementation, in Functional Electrodes for Enzymatic and Microbial Bioelectrochemical Systems, V. Flexer and N. Brun, Editors. 2017, World Scientific.

34. Patil, S.A., S. Gildemyn, D. Pant, K. Zengler, B.E. Logan, and K. Rabaey, A logical data representation framework for electricity-driven bioproduction processes. Biotechnology Advances, 2015. 33(6): p. 736–744.

35. Romans-Casas, M., L. Feliu-Paradeda, M. Tedesco, H.V.M. Hamelers, L. Bañeras, M.D. Balaguer, S. Puig, and P. Dessì, Selective butyric acid production from CO2 and its upgrade to butanol in microbial electrosynthesis cells. Environmental Science and Ecotechnology, 2023: p. 100303.

36. Zhang, F., J. Ding, Y. Zhang, M. Chen, Z.-W. Ding, M.C.M. van Loosdrecht, and R.J. Zeng, Fatty acids production from hydrogen and carbon dioxide by mixed culture in the membrane biofilm reactor. Water Research, 2013. 47(16): p. 6122–6129.

37. Ammam, F., P.-L. Tremblay, D.M. Lizak, and T. Zhang, Effect of tungstate on acetate and ethanol production by the electrosynthetic bacterium Sporomusa ovata. Biotechnology for Biofuels, 2016. 9(1): p. 163.

38. Roghair, M., D.P.B.T.B. Strik, K.J.J. Steinbusch, R.A. Weusthuis, M.E. Bruins, and C.J.N. Buisman, Granular sludge formation and characterization in a chain elongation process. Process Biochemistry, 2016. 51(10): p. 1594–1598.

39. Wang, D., Q. Liang, N. Chu, R.J. Zeng, and Y. Jiang, Deciphering mixotrophic microbial electrosynthesis with shifting product spectrum by genome-centric metagenomics. Chemical Engineering Journal, 2023. 451: p. 139010.

40. Marshall, C., D. Ross, K. Handley, P. Weisenhorn, J. Edirisinghe, C. Henry, J. Gilbert, H. May, and R.S. Norman, Metabolic reconstruction and modeling microbial electrosynthesis. BioRxiv, 2017: p. 059410.

41. Ross, D.E., C.W. Marshall, H.D. May, and R.S. Norman, Metagenome-Assembled Genome Sequences of Acetobacterium sp. Strain MES1 and Desulfovibrio sp. Strain MES5 from a Cathode-Associated Acetogenic Microbial Community. Genome Announcements, 2017. 5(36).

42. Bengelsdorf, F.R., M.H. Beck, C. Erz, S. Hoffmeister, M.M. Karl, P. Riegler, S. Wirth, A. Poehlein, D. Weuster-Botz, and P. Dürre, Chapter Four - Bacterial Anaerobic Synthesis Gas (Syngas) and CO2+H2 Fermentation, in Advances in Applied Microbiology, S. Sariaslani and G.M. Gadd, Editors. 2018, Academic Press. p. 143–221.

43. Litty, D. and V. Müller, Butyrate production in the acetogen Eubacterium limosum is dependent on the carbon and energy source. Microbial Biotechnology, 2021. 14(6): p. 2686–2692.

44. Lan, P.T.N., M. Sakamoto, S. Sakata, and Y. Benno, Bacteroides barnesiae sp. nov., Bacteroides salanitronis sp. nov. and Bacteroides gallinarum sp. nov., isolated from chicken caecum. International Journal of Systematic and Evolutionary Microbiology, 2006. 56(12): p. 2853–2859.

45. Iino, T., K. Mori, K. Tanaka, K.-i. Suzuki, and S. Harayama, Oscillibacter valericigenes gen. nov., sp. nov., a valerate-producing anaerobic bacterium isolated from the alimentary canal of a Japanese corbicula clam. International journal of systematic and evolutionary microbiology, 2007. 57(8): p. 1840–1845.

46. Raes, S.M.T., L. Jourdin, C.J.N. Buisman, and D.P.B.T.B. Strik, Continuous Long-Term Bioelectrochemical Chain Elongation to Butyrate. ChemElectroChem, 2017. 4(2): p. 386–395.

