## Supplemental information for "Microbial Electrosynthesis from CO_2_ reaches Productivity of Syngas and Chain Elongation Fermentations"

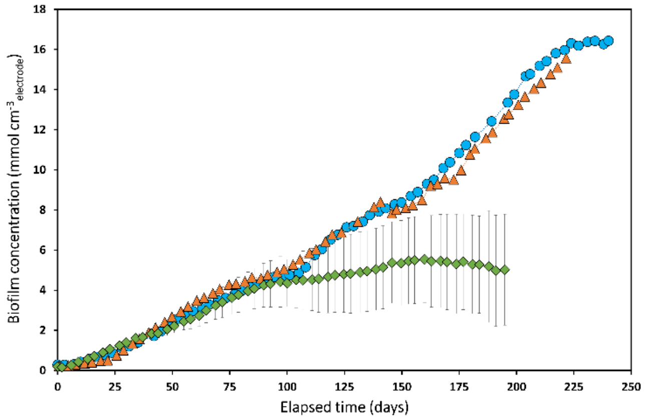


**Figure S1 Biofilm concentration measured over time in the galvanostatic (orange triangles) and potentiostatic (blue circles) reactors, compared with the state-of-the-art from Winkelhorst et al. 2023 (green diamonds).**

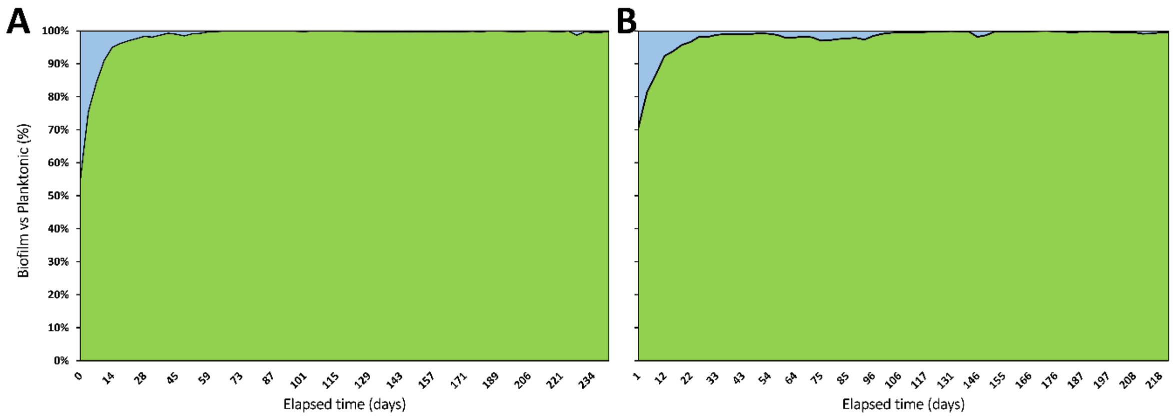


**Figure S2 Relative abundance of biomass in biofilm and in suspension in the galvanostatic (A) and potentiostatic (B) reactors.**

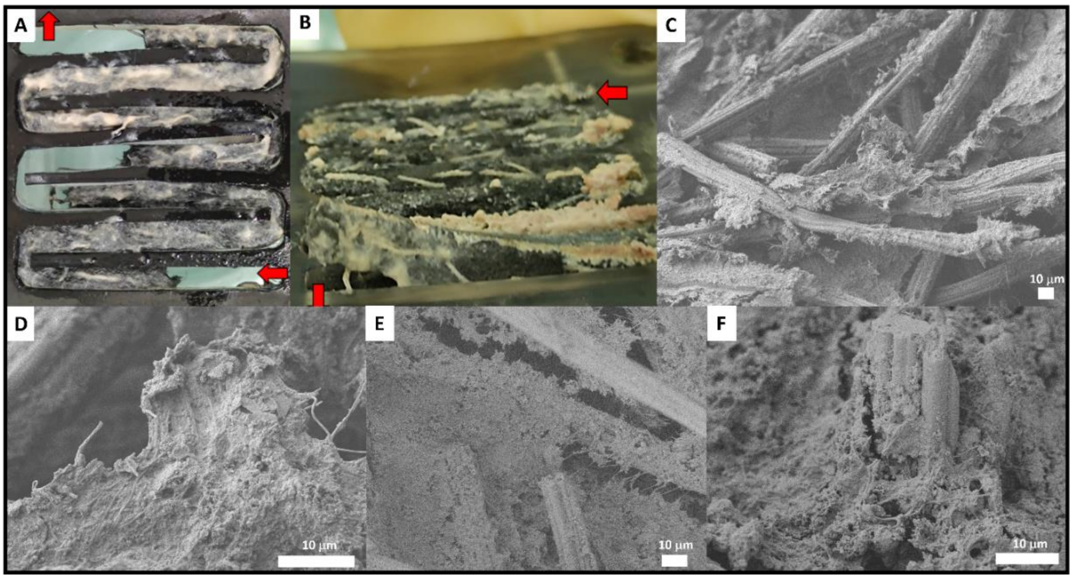


**Figure S3 Photograph of the biofilm on the side of the carbon felt facing the current collector (A) and throughout its thickness (B). SEM images of the biofilm grown on the carbon felt electrode of the potentiostatic (C-D) and galvanostatic (E-F) reactors, at the end of the experiment.**

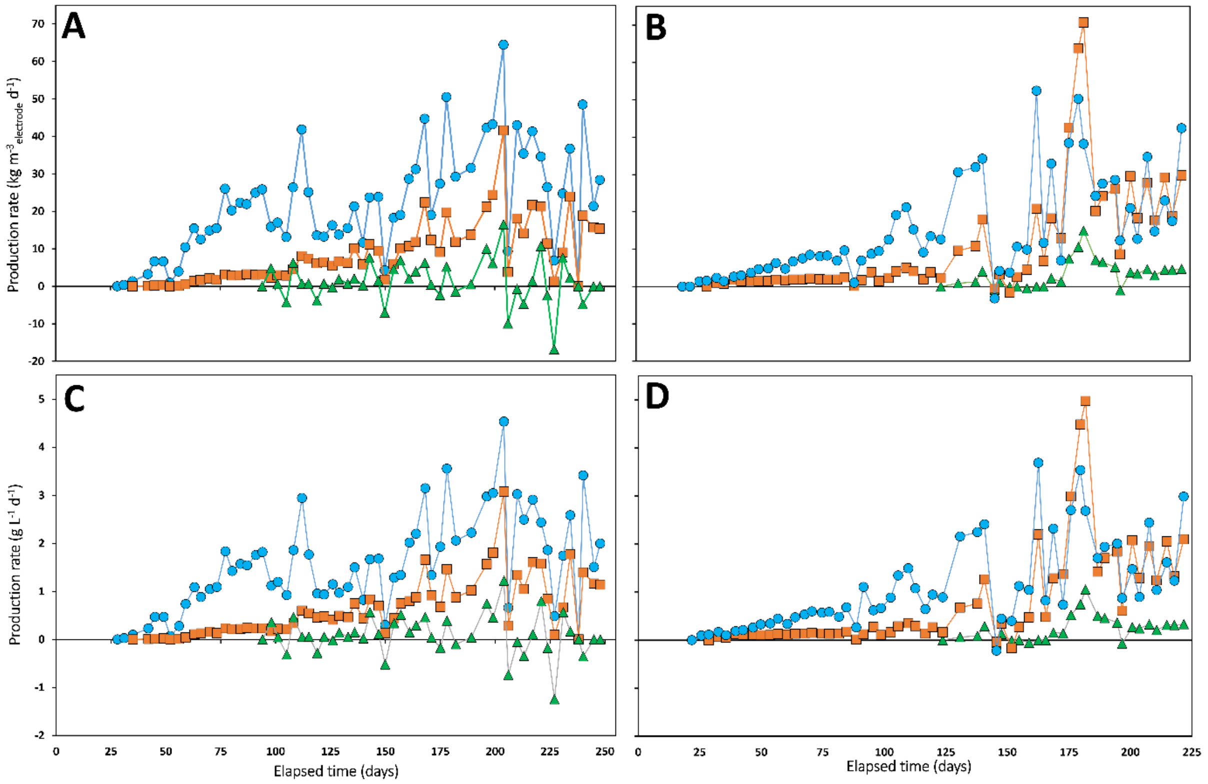


**Figure S4 Production rates, normalized to electrode volume (A-B) and total catholyte volume (C-D), of acetate (blue circles), butyrate (orange squares) and caproate (green triangles) measured over time in the galvanostatic (A, C) and potentiostatic (B, D) reactors.**

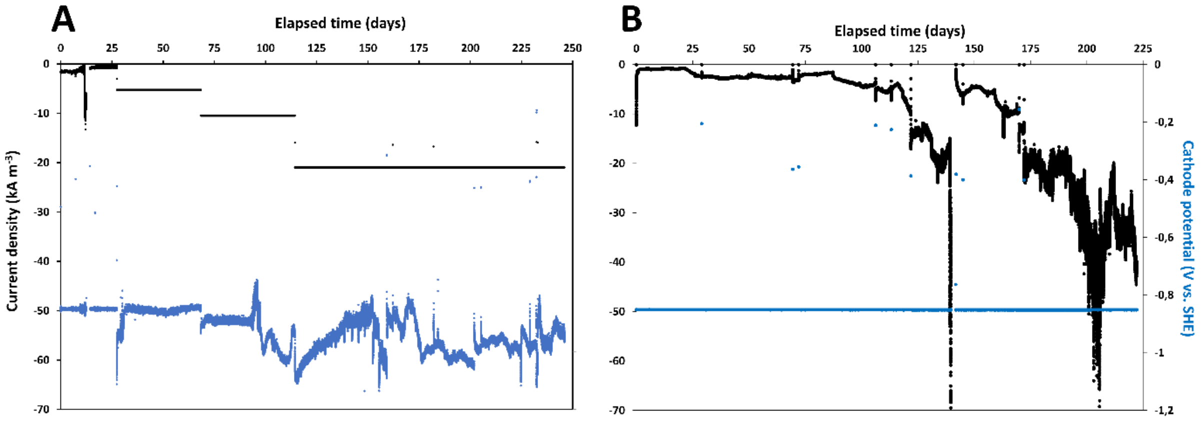


**Figure S5 Current profile in kA $m^{-3}_{cathode}$ in the galvanostatic (A) and potentiostatic (B) reactors.**

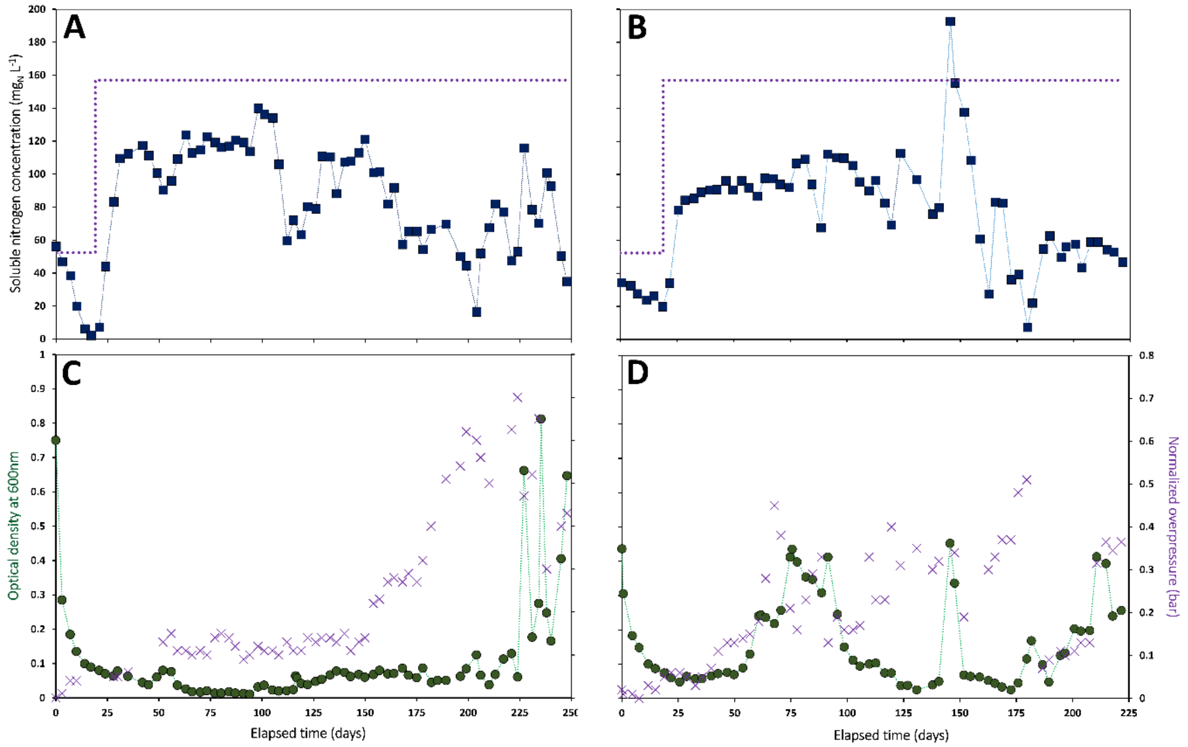


**Figure S6 Soluble nitrogen concentration (A-B), optical density and normalized overpressure (C-D) measured over time in the galvanostatic (A & C) and potentiostatic (B & D) reactors.**

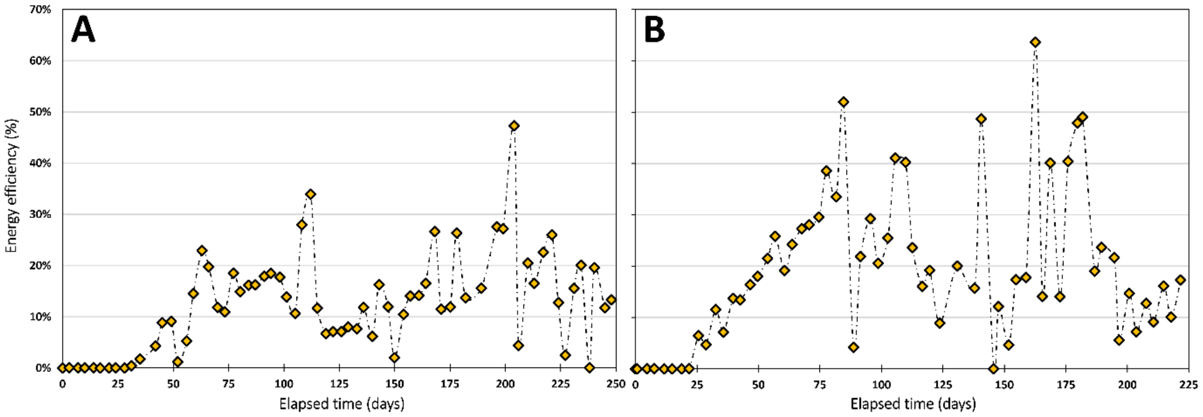


**Figure S7 Energy efficiency over time in the galvanostatic (A) and potentiostatic (B) reactors.**

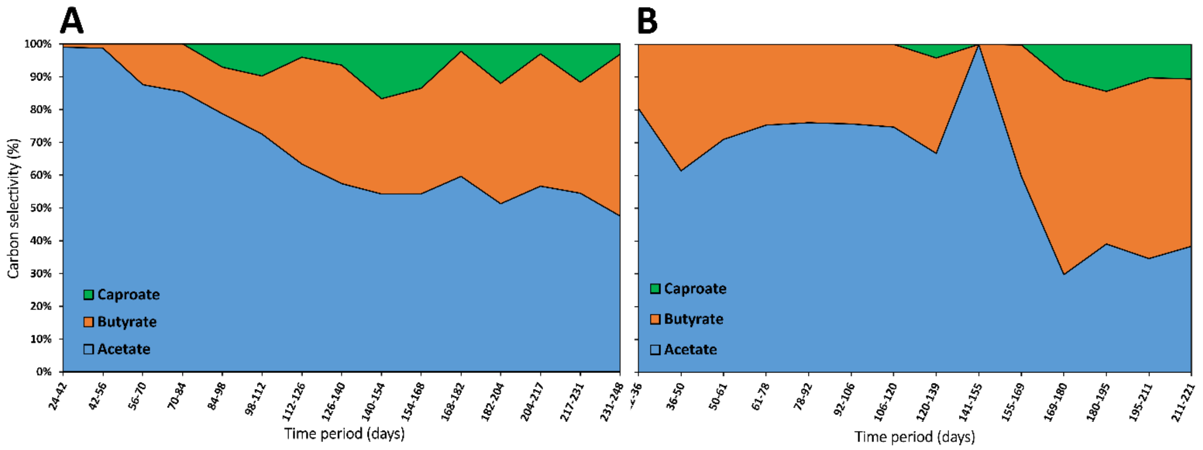


**Figure S8 Carbon selectivity over time in the galvanostatic (A) and potentiostatic (B) reactors.**

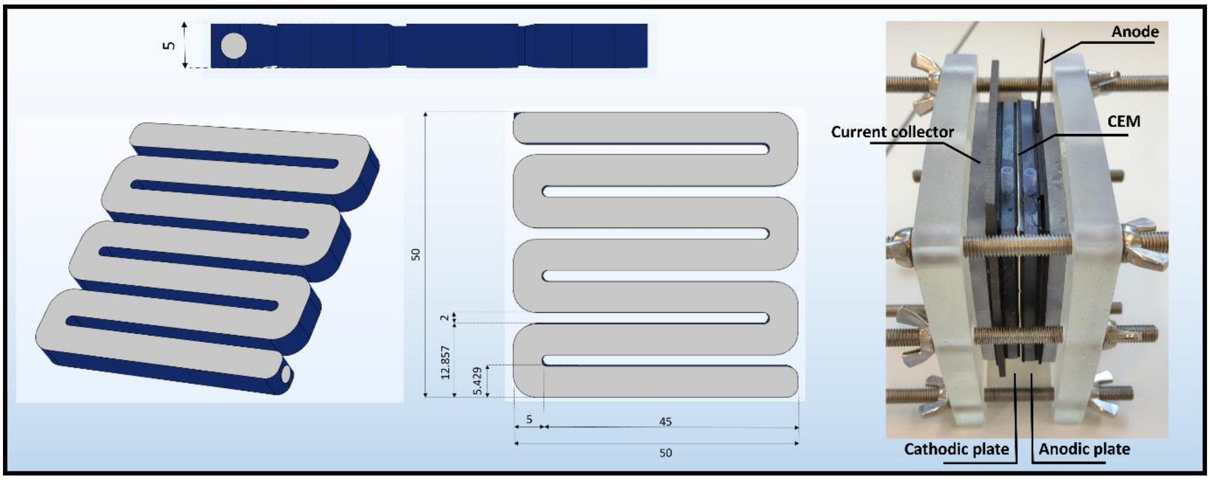


**Figure S9 Reactor scheme with dimensions (left and middle) and photo of the reactor (right).**

**Table S1 Literature data used to make Figure 5B.**

| | | $CO_2 + H_2O$ + electricity (MES) | Syngas (CO, $CO_2$, $H_2$) fermentation | | |
|---|---|---|---|---|---|
| | | This study | Lee et al. 2019 - Lab-scale | Hefferman et al. 2020 - Lab-scale | Fackler et al. 2021 - LanzaTech |
| $q_P$ | $mol_C\ mol^{-1}\ d^{-1}$ | 0.2 | 9.5 | 12.4 | 19.9 |
| | % relative | 0.0 | 0.5 | 0.6 | 1.0 |
| Biomass concentration | $g_X\ L^{-1}$ | 390 | 0.5 | 0.5 | 2.5 |
| | % relative | 1 | 0.001 | 0.001 | 0.006 |
| Volumetric productivity | $g_C\ L^{-1}\ h^{-1}$ | 0.2 | 0.1 | 0.2 | 1.0 |
| | % relative | 0.2 | 0.1 | 0.2 | 1.0 |

### Sequencing data analysis

#### 1. Paired-end reads assembly and quality control

##### *1.1 Data split*

Paired-end reads was assigned to samples based on their unique barcode and truncated by cutting off the barcode and primer sequence.

##### *1.2 Sequence assembly*

Paired-end reads were merged using FLASH

(V1.2.7,http://ccb.jhu.edu/software/FLASH/) [1],a very fast and accurate analysis tool, which was designed to merge paired-end reads when at least some of the reads overlap the read generated from the opposite end of the same DNA fragment, and the splicing sequences were called raw tags.

##### *1.3 Data Filtration*

Quality filtering on the raw tags were performed under specific filtering conditions to obtain the high-quality clean tags[2]according to the QIIME(V1.7.0, http://qiime.org/index.html)[3]quality controlled process.

##### *1.4 Chimera removal*

The tags were compared with the reference database(Gold database, http://drive5.com/uchime/uchime_download.html)using UCHIME algorithm(UCHIME Algorithm, http://www.drive5.com/usearch/manual/uchime_algo.html)[4]to detect chimera sequences, and then the chimera sequences were removed[5]. Then the Effective Tags finally obtained.

#### 2. ASV Denoise and Species Annotation

##### *2.1 ASVs Denoise*

For the Effective Tags obtained previously, denoise was performed with DADA2 or deblur module in the QIIME2 software (Version QIIME2-202006) to obtain initial ASVs (Amplicon Sequence Variants) (default: DADA2), and then ASVs with abundance less than 5 were filtered out [6].

##### *2.2 Species annotation*

Species annotation was performed using QIIME2 software. For 16S/18S, the annotation database is Silva Database, while for ITS, it is Unite Database.

##### *2.3 Phylogenetic relationship Construction*

In order to study phylogenetic relationship of each ASV and the differences of the dominant species among different samples(groups), multiple sequence alignment was performed using QIIME2 software.

##### *2.4 Data Normalization*

The absolute abundance of ASVs was normalized using a standard of sequence number corresponding to the sample with the least sequences. Subsequent analysis of alpha diversity and beta diversity were all performed based on the output normalized data.

#### 3. Alpha Diversity

In order to analyze the diversity, richness and uniformity of the communities in the sample, alpha diversity was calculated from 7 indices in QIIME2, including Observed_otus, Chao1, Shannon, Simpson, Dominance, Good's coverage and Pielou_e.

Three indices were selected to identify community richness:
Observed_otus – the number of observed species (http://scikit-bio.org/docs/latest/generated/skbio.diversity.alpha.observed_otus.html);

Chao – the Chao1 estimator (http://scikit-bio.org/docs/latest/generated/skbio.diversity.alpha.chao1.html);
Dominance – the Dominance index (http://scikit-bio.org/docs/latest/generated/skbio.diversity.alpha.dominance.html);
Two indices were used to identify community diversity:
Shannon – the Shannon index (http://scikit-bio.org/docs/latest/generated/skbio.diversity.alpha.shannon.html);
Simpson – the Simpson index (http://scikit-bio.org/docs/latest/generated/skbio.diversity.alpha.simpson.html);
One indice was used to calculate sequencing depth:
Coverage – the Good's coverage (http://scikit-bio.org/docs/latest/generated/skbio.diversity.alpha.goods_coverage.html);
One indice was used to calculate species evenness:
Pielou_e – Pielou's evenness index (http://scikit-bio.org/docs/latest/generated/skbio.diversity.alpha.pielou_e.html).

#### 4. Beta Diversity

In order to evaluate the complexity of the community composition and compare the differences between samples(groups), beta diversity was calculated based on weighted and unweighted unifrac distances in QIIME2.

Cluster analysis was performed with principal component analysis (PCA), which was applied to reduce the dimension of the original variables using the ade4 package and ggplot2 package in R software (Version 3.5.3).

Principal Coordinate Analysis (PCoA) was performed to obtain principal coordinates and visualize differences of samples in complex multi-dimensional data. A matrix of weighted or unweighted unifrac distances among samples obtained previously was transformed into a new set of orthogonal axes, where the maximum variation factor was demonstrated by the first principal coordinate, and the second maximum variation factor was demonstrated by the second principal coordinate, and so on. The three-dimensional PCoA results were displayed using QIIME2 package,

while the two-dimensional PCoA results were displayed using ade4 package and ggplot2 package in R software (Version 2.15.3).

To study the significance of the differences in community structure between groups, the adonis and anosim functions in the QIIME2 software were used to do analysis. To find out the significantly different species at each taxonomic level (Phylum, Class, Order, Family, Genus, Species), the R software (Version 3.5.3) was used to do MetaStat and T-test analysis. The LEfSe software (Version 1.0) was used to do LEfSe analysis (LDA score threshold: 4) so as to find out the biomarkers. Further, to study the functions of the communities in the samples and find out the different functions of the communities in the different groups, the PICRUSt2 software (Version 2.1.2-b) was used for function annotation analysis.

References

[1] Magoč T, Salzberg S L. FLASH: fast length adjustment of short reads to improve genome assemblies. Bioinformatics 27.21 (2011): 2957-2963.

[2] Bokulich, Nicholas A., et al. Quality-filtering vastly improves diversity estimates from Illuminaamplicon sequencing. Nature methods 10.1 (2013): 57-59.

[3] Caporaso, J. Gregory, et al. QIIME allows analysis of high-throughput community sequencing data. Nature methods 7.5 (2010): 335-336.

[4] Edgar, Robert C., et al. UCHIME improves sensitivity and speed of chimera detection. Bioinformatics 27.16 (2011): 2194-2200.

[5] Haas, Brian J., et al. Chimeric 16S rRNA sequence formation and detection in Sanger and 454-pyrosequenced PCR amplicons.Genome research 21.3 (2011):494-504.

[6] Wang, Qiong, et al. Naive Bayesian classifier for rapid assignment of rRNA sequences into the new bacterial taxonomy. Applied and environmental microbiology 73.16 (2007): 5261-5267.
